# Fatigue does not increase limb asymmetry or induce proximal joint power shift during sprinting in habitual, multi-speed runners

**DOI:** 10.1101/2021.10.17.464459

**Authors:** Shayne Vial, Jodie Cochrane Wilkie, Mitchell Anthony, Mitchell Turner, J. Blazevich

## Abstract

The ability to shift from walking and jogging to sprinting gaits, even when fatigued after prolonged effort, would have been as useful to our hunter-gatherer ancestors as it is to modern athletes. During prolonged jogging, joint moment and work decrease in the distal (ankle) joint but increase at proximal (hip/knee) joints as fatigue progresses, and such adaptations might be expected to occur in sprinting. Fatigue is also thought to increase inter-limb kinematic and force production asymmetries, which are speculated to influence injury risk. However, the effects of running-related fatigue on sprint running gait have been incompletely studied, so these hypotheses remain untested. We studied 3-D kinematics and ground reaction force production in dominant (DL) and non-dominant (NDL) legs during both non-fatigued and fatigued sprinting in habitual but uncoached running athletes. Contrary to the tested hypotheses, relative between-leg differences were greater in non-fatigued than fatigued sprinting. When not fatigued, DL produced greater propulsive impulse through both greater positive and negative work being done at the ankle, whilst NDL produced more vertical impulse, possibly resulting from the greater hip flexion observed prior to the downwards acceleration of the foot towards the ground. Whilst few changes were detected in DL once fatigued, NDL shifted towards greater horizontal force production, largely resulting from an increase in plantarflexion (distal-joint) moments and power. After fatiguing running, therefore, inter-limb asymmetry was reduced during sprinting and no distal-to-proximal shift in work/power was detected. Speculatively, these adaptations may help to attenuate decreases in running speed whilst minimising injury risk.

**Significance:** The ability to attain fast running speeds may critically determine success in tasks such as prey chase-and-capture in hunter-gatherer societies as well as success in modern sports competitions. At times, sprint running may have to be performed whilst fatigued from previous, longer-distance running, when speeds are reduced, and injury risk may be higher. Previous work indicated that fatigue prompts a proximal shift in joint work and power production and an increase in inter-limb asymmetry. On the contrary, we show that relative ankle positive and negative joint work was maintained in the face of fatigue and that inter-limb asymmetry was reduced in a group of runners experienced, but not formally instructed, in both long-distance and sprint running.

## Introduction

Anatomical adaptations present in *Homo erectus*, including relatively long legs, short arms, wide shoulders, and narrow waist, were critical to our successful transition towards terrestrial bipedality (Lieberman et al., 2009; Steudel-Numbers et al., 2007). These adaptations, which are even more prominent in modern humans, would have benefited migration and enhanced our persistent scavenging and hunting capacities (3, 4), and thus contributed to our success as a species. In addition to walking, running would have been relatively common as it played a part in allowing humans to track and capture animals over long distances as well as reducing exposure time to both predators and the environment (5, 6). Today, running is not only a common pastime but an integral part of many games and sports. Consistent with its importance, researchers have developed a detailed biomechanical picture of human running under varying environmental conditions in both pre-industrial, non-sports-trained (assumedly similar to hunter-gatherer) (7–9) and post-industrial, sports-trained (10–14) populations.

Nonetheless, prior to industrialisation, substantial changes in migration pace would also have been needed, including slowing when prey tracking was difficult or the animal reduced speed then speeding up or sprinting as the target raced or was in need of startling to prompt the continued movement needed to tire it (15). Bursts of high-speed running would also have been necessary when confronted by a predator, either human or other animal. Early humans, therefore, would have needed to run at a range of speeds for varying durations, shifting between walking, endurance running and, to a lesser extent, sprinting (3). And, perhaps reflective of its historical importance, this capacity is also challenged in many of our modern games and sports.

Because a shift from walking or jogging to sprinting may have been needed after covering great distances, sprint efforts would have, at least sometimes, been performed whilst in a state of fatigue (4, 15). Such fatigue decreases maximal force and power capacity during continued exercise (16–18) and negatively impacts sprinting speed (19–21). Although in most modern societies there is now no requirement for long tracking or hunting efforts, endurance running interspersed with sprinting is still exceptionally common in sports (22–25). These sprints are typically performed during fatigue even though we also tend to reduce the number of voluntary sprint efforts in the latter stages of a match or after periods of high exertion (Mohr et al., 2004; Mohr et al., 2005; Reilly et al., 2008). Despite this, comparatively little is known about how we run at high speeds whilst fatigued, so it is not known whether we substantively alter running technique in an attempt to maintain sprinting speed despite, or as a direct result of, this fatigue.

Fatigue-related gait alterations may be expected in response to decreased muscle force or power, but such alterations might also be causative of the decrements in force and power through changes in the muscle lengths adopted and muscle shortening speeds reached when the gait pattern is altered (28–30). Reduced work performed at the ankle, for example, has been observed with concomitant increased joint work about the knee and hip (i.e. a disto-proximal shift) during both fatiguing, persistent jogging (31), and higher-speed runs (e.g. ~180 s to exhaustion; Willer et al., 2021). However, it is not known whether such a shift from highly elastic-powered ankle propulsion to muscle-dominant knee-hip power production also occurs during maximal effort sprinting after fatiguing running exercise.

Additionally, human gait is usually performed relatively symmetrically between limbs, although small inter-limb differences exist (32–34). Nonetheless, stronger, less fatigable muscles might be expected to be recruited more whilst weaker, fatigued muscles might be rested or at least provide less power, during fatigued sprinting, even when a high overall power output (running speed) is required. Given that humans show significant lateral bias, i.e. handedness (35), this might conceivably lead to power production asymmetries during running (36–38). Asymmetries in muscle force production measured in strength tests have been associated with increases in injury risk (Lord et al., 2018; Stephens et al., 2005), leading to speculation that substantive force or power production asymmetries may be inherently injurious (32, 41). It is also not known whether technique adaptations might influence injury risk, which would have been problematic for our ancestors during hunting or escape and is particularly problematic for modern competitive athletes. Whether gait asymmetry is directly causative of injury, or even whether substantive asymmetries evolve during sprinting after prior bouts of fatiguing running, remains unclear (42–44). Hence, it is not known whether fatigue-induced gait asymmetries might have been a problem for our ancestors or, indeed, are a current problem for modern running-based athletes.

Here, we comprehensively explore kinematic and kinetic patterns of dominant (strong) and non-dominant (weak) legs during both non-fatigued and fatigued maximal sprinting in order to describe differences in the response of each leg to fatiguing running exercise. We tested the hypothesis that substantial acute kinematic and kinetic adaptations would be observed during fatigue, and that generally small inter-limb asymmetries would be exacerbated, in a group of highly experienced runners.

## Methods

Thirteen semi-professional male Association Football players (age: 19.1 ± 2.1 y, body mass: 72.5 ± 6.9 kg, height: 175 ± 7.7 cm) volunteered for the study. The athletes regularly performed sprint running as well as lower intensity endurance running both in competitive (i.e. stressful) games as well as in their (less stressful) training but had not received any formal running technique instruction. Thus, they present a cohort who have a freely chosen running method. They also performed no formal strength or other supplementary training that might influence running performance or their response to fatiguing running exercise. Additional rationale for the choice of study cohort (e.g. over participants in track & field or other sports) as well as participant data are presented in Supplementary Information. All subjects were free from injury for at least 6 months before testing, wore their normal training attire, and wore the same (their own) running shoes during testing. The study was approved by Edith Cowan University of Human Ethics Committee and players gave written consent prior to testing.

### Biomechanical measurements

On arrival, height and body mass were recorded for each subject and then a custom-defined set of retroreflective cluster-based markers used for the 3D motion analysis were firmly but comfortably attached to identified anatomical landmarks (Table S1). The retroreflective markers were captured during the trials by thirteen VICON motion analysis cameras (Oxford Metrics Ltd., Oxford, UK) set at a frame rate of 250 Hz. Motion data capture was synchronised with ground reaction force data, recorded with five serially arranged 600 x 900-mm in-ground triaxial force platforms (Kistler Quattro, Type9290AD, Victoria, Australia) at an analogue-digital conversation rate of 1000 Hz. Motion capture cameras were positioned to ensure a suitable capture volume around the in-ground force platforms to capture the sprint running trials (Figure S1). In order for the anatomical markers to be referenced to the tracking markers prior to recording sprint running trials, static subject calibration trials were obtained with the subject standing in the anatomical position, following by dynamic calibration trials in which subjects moved their legs through a range of motion to enable post-collection determination of functional joint centres using Visual 3D software (C-Motion, Germantown, MD, USA).

### Protocol

After completing the static and dynamic calibration trials, subjects performed a comprehensive standardised warm-up (~15-20 min) at their chosen intensity (Table S2). Then, three single-leg vertical jumps (SLVJ) were performed on each leg to obtain jump height (Figure S2) as a measure to determine the dominant and non-dominant limb (the SLVJ requires significant force production, skill, and coordination – see section 5 in Supplementary Information). In some studies, researchers have designated the preferred kicking leg as dominant, which may not be the stronger of the two legs and thus may not accord with our definition. After SLVJs, three maximal 50-m sprint running efforts were performed from a standing start. The starting point was located 40-m from the last force platform in the series to enable force recordings during the maximal velocity phase (i.e. 35-40 m). The end point of the sprint was located 50 m from the start point to ensure subjects did not decelerate through the data capture zone. The subjects were allowed exactly 60 s of rest between trials; the relatively short rest was used to minimise the recovery from fatigue in post-running trials but did not induce detectable running fatigue in the non-fatigued tests (see Results).

After the first set of sprints (pre-fatigue test), subjects completed a soccer-specific fatiguing exercise protocol (Ball–Sport Endurance and Sprint Test; BEAST 45, Figure S3) lasting 45 min (45) that included repeated bouts of sprint running, jogging, changes of direction, walking, backward running, and stationary recovery similar to the first half of a soccer match. This protocol was chosen as it was familiar to the subjects, who could therefore complete it without a notable pacing strategy or extensive familiarisation, and because it incorporated all directions of movement that might be done in either a hunt or the chase of an agile animal as well as in modern sports. After performing the fatiguing exercise, subjects jogged to the start line in ~80 s. Once at the start line, a countdown from 5 to 1 led into the first sprint effort. The subjects performed three further maximal sprinting trials (post-fatigue test) with 60-s inter-sprint rests.

### Data analysis

All trials were digitised using VICON Nexus software (Oxford Metrics Ltd., Oxford, UK). Trials were then exported as C3D files to Visual 3D (C-Motion, Germantown, MD, USA) and both ground reaction force and marker trajectory data were filtered using a fourth-order (zero lag) low-pass Butterworth filter with a 15-Hz cut-off frequency, determined by residual analysis (46, 47). Using Visual 3D software, static calibration data, subject height, and body mass were used to create an individually scaled skeletal model that included the trunk, pelvis, thigh, shank, and foot segments using standard available inertial parameters (segment mass (48) and moments of inertia (49)) in Visual 3D. For both limbs in the sprint running trials, sagittal segment angles were calculated relative to the laboratory reference frame and were normalised to normal upright standing position, while conventional Visual 3D (C-Motion) calculation methods using Newton-Euler procedures were used to compute joint moments and powers during the second half of retraction, defined as the forward rotation of the leg up to peak hip flexion, and protraction, defined as the backward rotation of the leg from peak hip flexion to toe off (see Figure 1).

**Figure 1.**
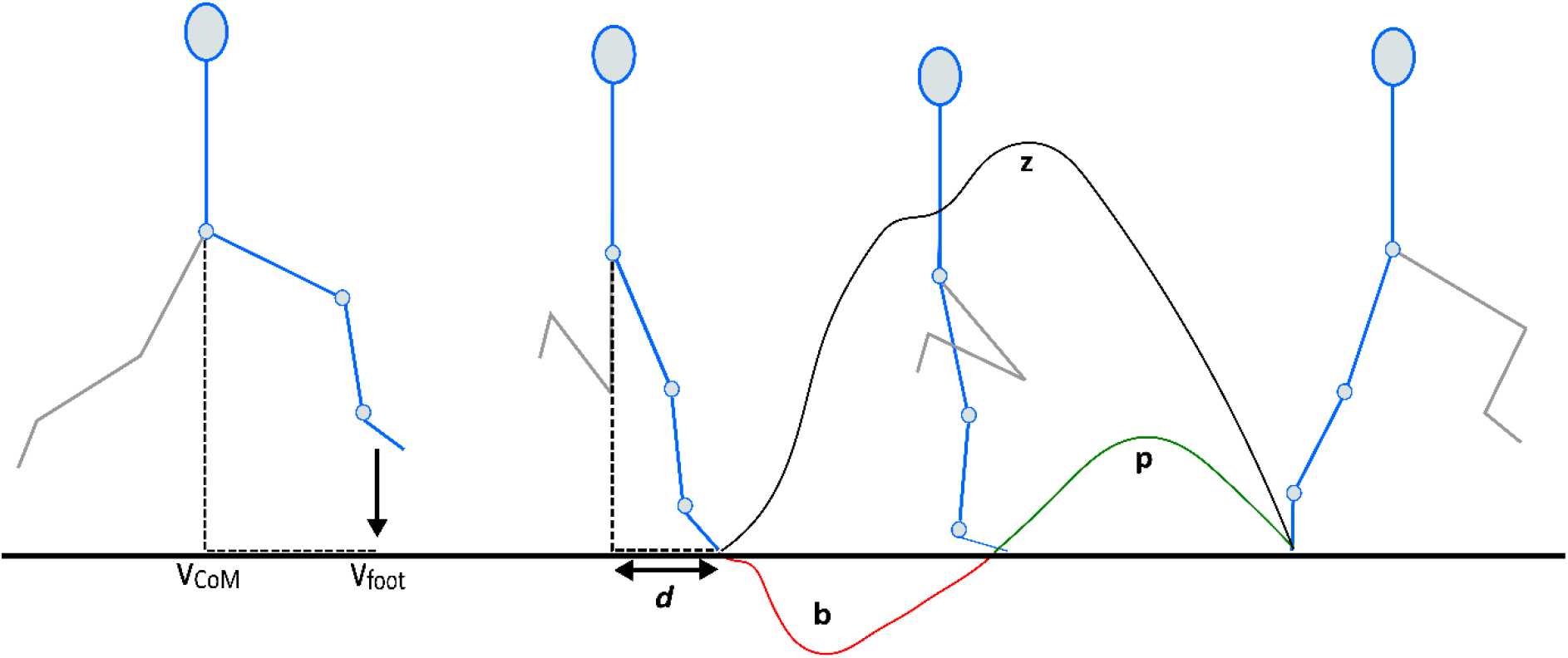
Second half of retraction and protraction phases. Key variables compared between dominant and non-dominant legs in non-fatigued and fatigued sprinting. Vertical centre of mass velocity (v_CoM_), vertical foot velocity (v_foot_), anterior-posterior distance of foot relative to CoM at foot-strike (*d*), braking impulse (b), propulsive impulse (p), vertical impulse (z).

### Statistical Analysis

Descriptive statistics (means and standard deviations) were obtained for variables captured in both the non-fatigued and fatigued conditions. For discrete variables, paired t-tests were used to compare between the dominant and non-dominant legs as well as non-fatigued and fatigued trials. Statistical Parametric Mapping (SPM) was used to compared joint kinematic and kinetic data during the second half of the retraction and protraction phase between the dominant and non-dominant legs across both conditions using open-source SPM code (SPM1D open-source package, spm1d.org) in Python (50). Joint kinematic and kinetic data were normalised, representing 0%-100% of the retraction and protraction phases (see Figure 1 below), respectively. An SPM paired t-test was then performed separately at each of the 101 time points resulting in the output of a statistical parametric map (SPM{t}). When the SPM{t} exceeded the critical threshold, the variable was considered significantly different between legs or conditions, and a collection of ≥5 consecutive points exceeding the threshold was considered statistically meaningful (51). All statistical analyses for discrete variables were performed using JAMOVI (Version 1.6, Sydney, Australia).

## Results

The key sprint performance variables compared between dominant (DL; stronger) and non-dominant (NDL; weaker) legs in non-fatigued and fatigued sprinting are shown in Figure 1. DL was selected as the leg that produced the greatest jump height in the single-leg vertical jump test, with non-fatigued DL jumps being 22.7 ± 0.9 cm and NDL jumps being 21.1 ±1.5 cm (Figure S2). The average maximum horizontal velocity of the CoM during non-fatigued sprinting was 8.59 m/s, and decreased 0.33 m/s during fatigued sprinting (Table S4).

### Dominant (DL) vs. non-dominant (NDL) legs: Non-fatigued sprinting

Vertical foot velocity (i.e. towards the ground) relative to the centre of mass (CoM) was significantly greater in NDL (~7%) than DL immediately before foot-ground contact (Figure 2), however no statistical differences were observed in horizontal foot velocity. The anterior-posterior position of the foot relative to the CoM at foot-strike was closer to the CoM in DL than NDL (Figure 2). Accordingly, a smaller braking impulse (~7%) and greater propulsive impulse (~14%), but smaller vertical impulse (~8%), was produced by DL than NDL (Table 1) with similar ground contact times (Figure S4).

**Figure 2.**
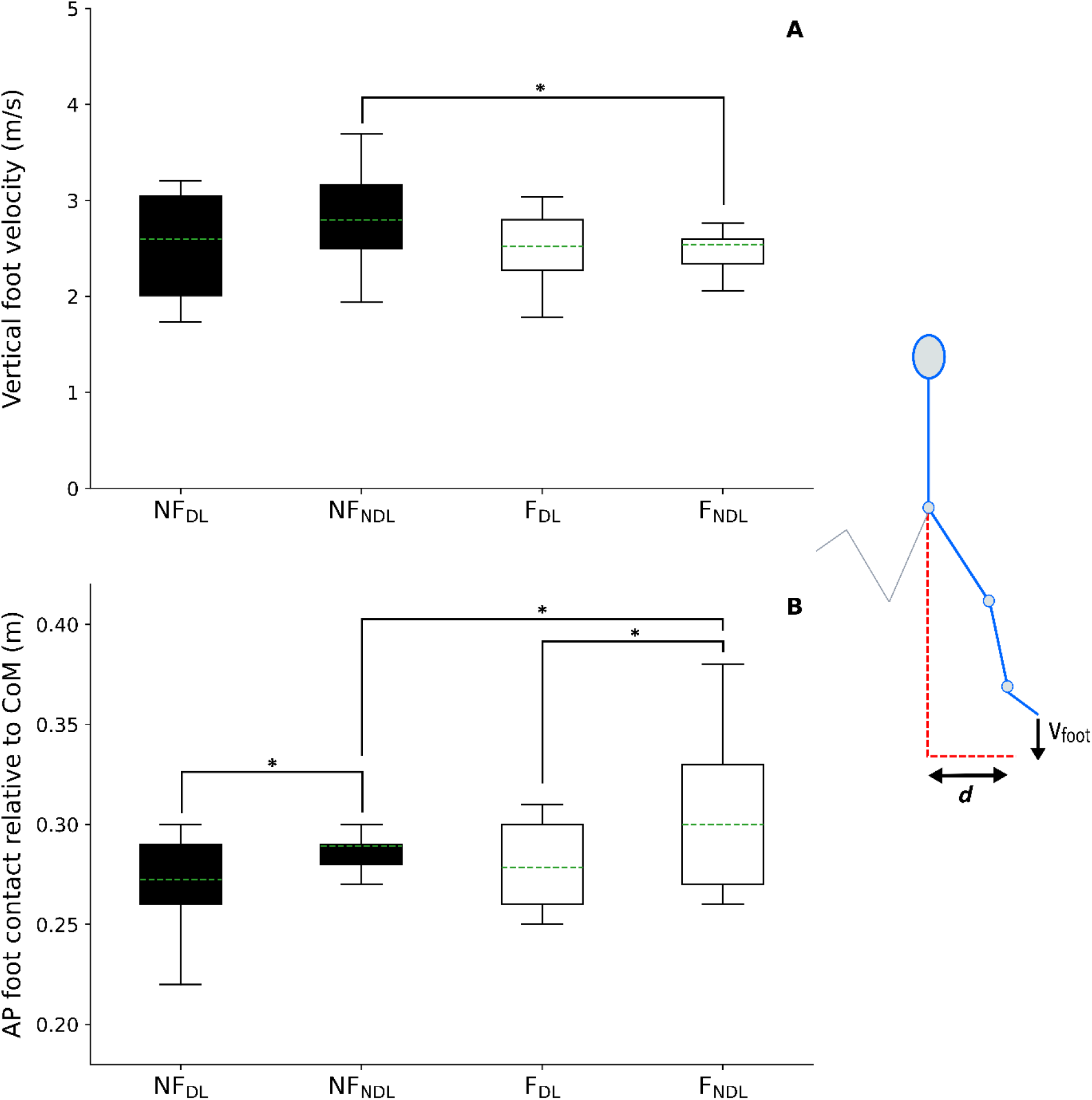
(A) Shows peak vertical velocity of the foot (metres per second) relative to the centre of mass velocity between early protraction to foot-strike; (B) anterior-posterior (AP) position of the foot at ground contact relative to the horizontal position of the centre of mass (metres) for the non-fatigued dominant leg (NF_DL_), non-fatigued non-dominant leg (NF_NDL_), fatigued dominant leg (F_DL_), and fatigued non-dominant leg (F_NDL_). V_foot_ represents the vertical velocity of the foot; *d* represents the anterior-posterior position of the foot relative to CoM at foot-strike. (A)* statistical difference of vertical velocity of the foot relative to CoM velocity, (B)* statistical difference of AP foot position relative to horizontal CoM position between legs and conditions, respectively (set at *p* < 0.05).

**Table 1.** Non-fatigued and fatigued braking, propulsive, vertical, and propulsive/braking impulse ratios for dominant and non-dominant legs. *statistical difference between dominant and non-dominant legs trials. ^statistical difference between non-fatigued and fatigued trials, respectively.

|  | Dominant leg | Non-dominant leg | Mean diff | 95% CI (change) |
| --- | --- | --- | --- | --- |
| | Mean ± SD | Mean ± SD | | |
| ***Non-fatigued*** |  |  |  |  |
| Braking impulse (kg·m/s) | -0.20 ± 0.10*^ | -0.21 ± 0.10* | -0.02 | -0.03, -0.02 |
| Propulsive impulse (kg·m/s) | 0.25 ± 0.10* | 0.21 ± 0.10*^ | 0.04 | -0.06, -0.02 |
| Vertical impulse (kg·m/s) | 2.30 ± 0.20* | 2.50 ± 0.20* | -0.20 | 0.10, 0.30 |
| Propulsive/braking ratio | 1.29 ± 0.10*^ | 1.04 ± 0.20* | -0.25 | -0.37, 0.13 |
| ***Fatigued*** |  |  |  |  |
| Braking impulse (kg·m/s) | -0.23 ± 0.10^ | -0.23 ± 0.10 | 0.07 | -0.02, 0.04 |
| Propulsive impulse (kg·m/s) | 0.24 ± 0.10 | 0.25 ± 0.02^ | -0.01 | -0.01, 0.02 |
| Vertical impulse (kg·m/s) | 2.40 ± 0.30 | 2.52 ± 0.20 | -0.10 | -0.07, 0.28 |
| Propulsive/braking ratio | 1.10 ± 0.20^ | 1.10 ± 0.20 | -0.03 | -0.04, 0.10 |

No differences were observed in sagittal plane pelvic kinematics between DL and NDL in the non-fatigued condition, however a greater peak hip flexion angle relative to the pelvis was observed in NDL during the retraction-protraction transition point (Table S5). The ankle was the dominant source of lower limb positive and negative joint work (J kg^−1^) for DL, whereas NDL had a relatively even distribution across the lower limb joints (Figure S5). Both limbs had the same positive (31%) and negative (34%) work performed at the hip joint. There was significantly more positive (~7%) and negative (~13%) work produced at the ankle in DL, whilst NDL produced more positive (~6%) and negative (~14%) work at the knee joint. In addition, a greater knee extension moment (Nm/kg) was observed from foot-strike through to early stance. By contrast, DL knee extension moment showed a more gradual increase and later peak than NDL (Figure 3) whilst the peak ankle plantar flexion moment was greater during the first half of stance in DL and a higher peak plantarflexor moment (~12%) was produced (Table S6).

**Figure 3.**
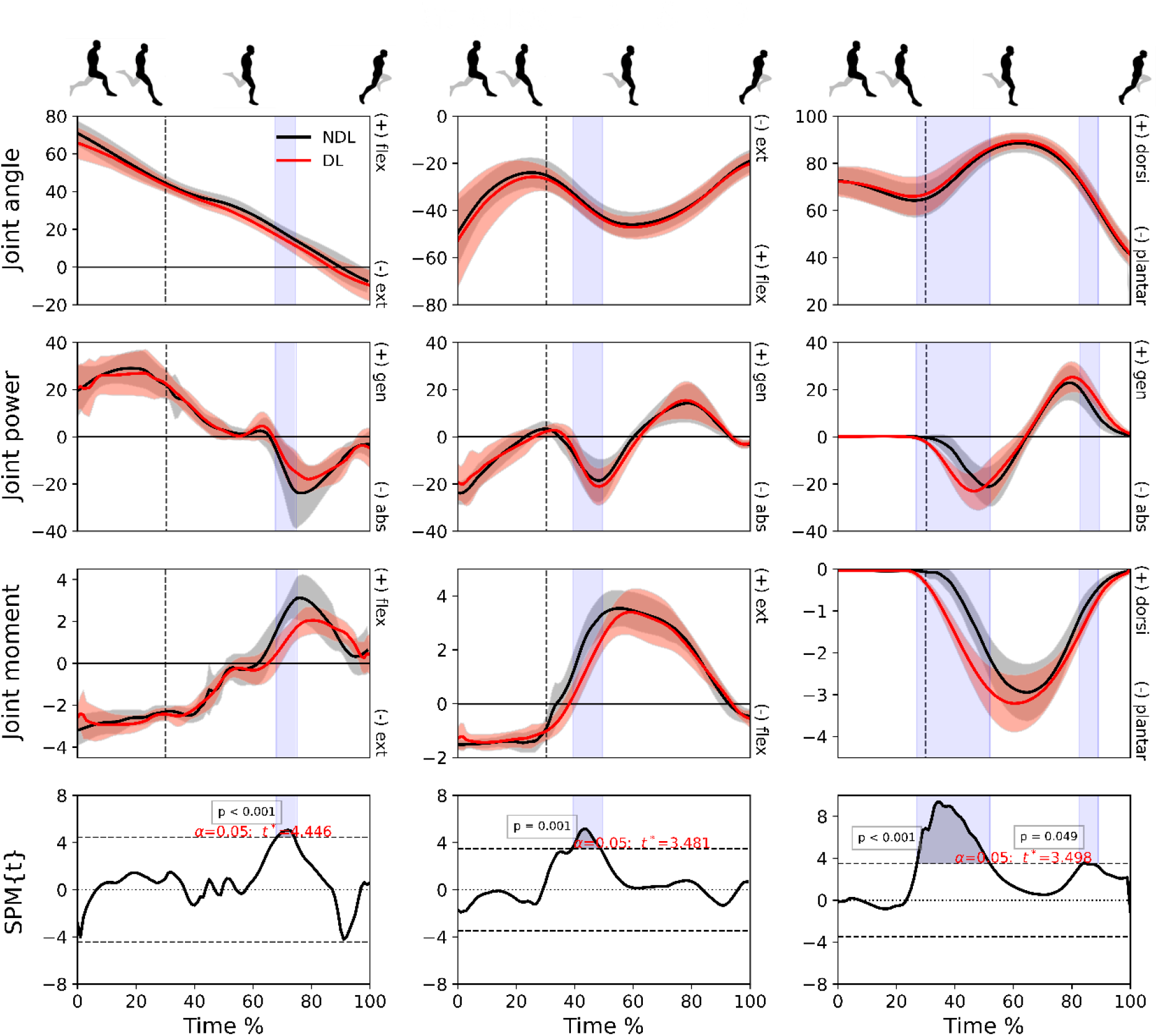
Mean (± standard deviation) joint angles (degrees), powers (W/kg) and moments (N/kg) at hip (column 1), knee (column 2) and ankle (column 3) joints for the dominant (red) and non-dominant (black) legs in the non-fatigued condition. Vertical dotted line represents foot-strike. The time-dependent paired t-values of the SPM (bottom row; set at *p* < 0.05) are shown as horizontal dashed lines. Shaded areas indicate regions with statistical differences.

### Effect of fatigue on DL leg and pelvis kinetics and kinematics

A decrease in positive (~7%) and no change in negative work (Figure S5) were observed at the hip joint whilst greater power was generated at the knee around mid-stance after the fatiguing running in DL, which corresponded with a greater braking impulse (~17%) (Table 1). After the fatiguing running exercise, the pelvis tilted more anteriorly (~3.6°) in the phase from early protraction phase to immediately prior to foot-ground contact when viewed from DL side (Table S5) whilst the SPM analyses revealed a significantly greater knee extension moment from foot-ground contact through to early stance (Figure S6).

### Effect of fatigue on NDL leg and pelvis kinetics and kinematics

During fatigued sprinting, in NDL, the ankle contributed slightly more positive and negative work (not statistically different) with similar hip and knee contributions (Figure S5). Furthermore, the pelvis tilted more anteriorly (~2.5°) between early protraction to immediately prior to foot-ground contact when viewed from NDL side (Table S5), and both peak hip flexion angle (~4%) and hip extension moment (~10%) decreased (Table S6). The knee remained more extended during early protraction in the fatigued condition but was similar by the point of foot-ground contact. Also, a much larger plantarflexion moment and greater power absorption was observed from foot-ground contact to early stance after the fatiguing exercise (Figure S7). This coincided with a reduced vertical foot velocity (Figure 2) just prior to foot-ground contact relative to the CoM yet greater propulsive impulse (~13%) (Table 1).

### Differential effects of fatigue on DL and NDL

Slightly more negative (~5%) work was performed at the knee in NDL than DL during fatigued sprinting (Figure S5). The SPM revealed a small difference in ankle joint kinetics (Figure 4) after the fatiguing running exercise. A slightly greater ankle plantarflexion moment was produced at foot-strike in DL than NDL and with shorter ground contact times (~3%), yet both legs produced similar braking, propulsive, and vertical impulses (Table 1). Less vertical CoM displacement (Figure S8) was observed during DL than NDL force production phases in the fatigued condition.

**Figure 4.**
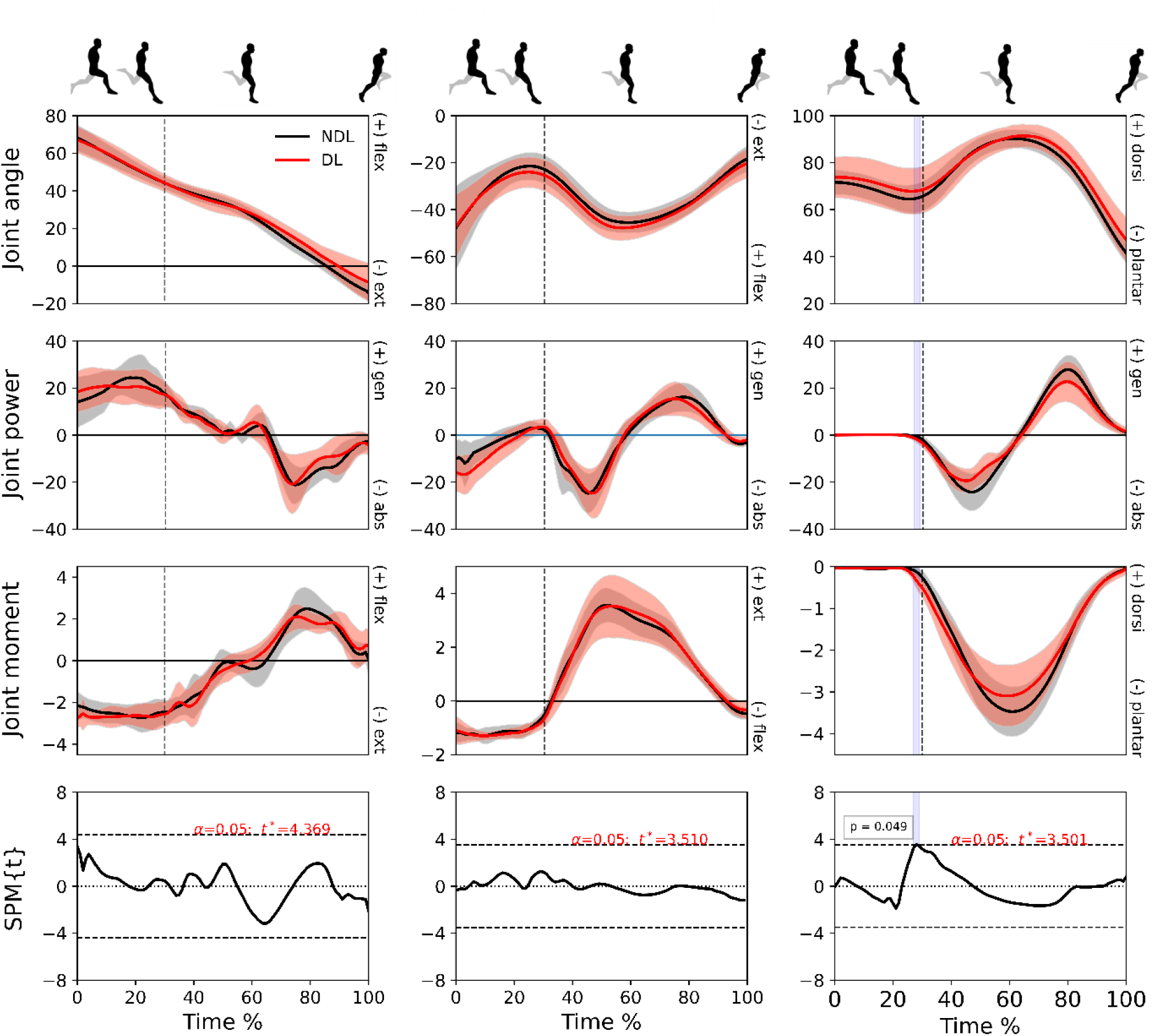
Mean (± standard deviation) joint angles (degrees), powers (W/kg) and moments (N/kg) at hip (column 1), knee (column 2) and ankle (column 3) joints for the dominant (red) and non-dominant (black) legs in the fatigued condition. Vertical dotted line represents foot-strike. The time-dependent paired t-values of the SPM (bottom row; set at *p* < 0.05) are shown as horizontal dashed lines. Shaded areas indicate regions with statistical differences.

See Figures S9-S12 in the Supplementary Information for visual representation of the differences observed between and within limbs and across conditions.

## Discussion

Contrary to the tested hypotheses, fatiguing running exercise completed by athletes who commonly practise both slower (endurance) and sprint-speed running did not promote a shift from elastic powered ankle propulsion to muscle-dominant knee-hip power production, and in fact a slightly earlier increase in plantarflexion moment was observed at ground contact in the weaker (non-dominant) limb in fatigued sprinting. Additionally, a reduction, rather than increase, in inter-limb asymmetry was observed, resulting from a decrease in the peak hip flexion angle at retraction-protraction transition and earlier rise in plantar flexion moment during foot contact in the weaker leg; after fatiguing running, no kinematic and few kinetic differences were detected between legs. Therefore, in our cohort of skilled runners, a strategy was adopted that maintained relative ankle positive and negative joint work in the face of fatigue and reduced inter-limb asymmetry. These acute adaptations to fatiguing running exercise contrast common hypotheses that are based on our understanding of acute adaptations during sub-maximal running (jogging) but may speculatively help to maintain high-speed power production whilst limiting asymmetry-induced injury risk during high-speed locomotion.

During non-fatigued sprinting, substantive differences in kinetic and kinematic variables were observed between the stronger (dominant leg; DL) and weaker (non-dominant leg; NDL) legs at the hip, knee, and ankle joints. DL produced greater propulsive impulse whereas NDL produced greater braking and vertical impulses during the ground-contact phase. Thus, the stronger leg tended to more critically influence forward velocity of the CoM whereas the weaker leg predominately projected the body with vertical velocity (assumedly to maintain stride length) with less assistance to forward propulsion (33, 38). A similar vertical CoM displacement was thus observed after projecting off either leg.

This inter-limb variability might be explained by technique differences between the legs. The peak hip flexion angle was greater in NDL at the retraction-protraction transition point (Table S5), without subsequent extensor moment increase. Accordingly, the vertical distance travelled by the foot to the ground and then the maximum vertical velocity of the foot both increased, with part of this effect potentially explained by ~24% greater hip extension angular velocity (although this varied individually and did not reach statistical significance; see Table S7). The greater vertical foot velocity would have better allowed it to move underneath the body in the short time available before ground contact to arrest the fall of the body’s CoM. Nonetheless, foot-strike still occurred further from the CoM in NDL than DL, contributing to a greater braking impulse and thus to a greater and earlier increase in knee extensor moment (i.e. a faster rate of joint moment development).

Conversely, in DL the ankle absorbed more power and the plantarflexion moment both commenced and reached an earlier and greater peak during ground contact. Therefore, joint moment production favoured knee extension in the weaker NDL but ankle plantarflexion in the stronger DL. If we assume that the energy stored in elastic tissues at each joint is linked to the work done (moment × angular displacement) in the early, i.e. “braking”, phase of ground contact, then the greater propulsive impulse provided by DL can be explained by use of a more ankle-dominant, energy storing strategy. Given these findings, one might conclude that the legs played different roles in, and thus contributed differently to, the sprint running cycle; whether this pattern maximises running speed remains to be explicitly tested in future studies.

Fatiguing running exercise reduced sprint running velocity (−4%) and increased ground contact times (~2% in DL and ~3% in NDL), and it slightly increased vertical CoM displacement and horizontal distance of the foot from the CoM at foot-ground contact in NDL. Nonetheless, when fatigued, both legs displayed similar peak hip flexion angles at the retraction-protraction transition point (in contrast to non-fatigued sprinting) and no differences were detected in knee or ankle joints ranges or in propulsive, braking, or vertical impulses. Furthermore, peak knee extensor moment increased in DL, so the difference to NDL that had been observed in non-fatigued sprinting was no longer present. Small but non-statistical increases in NDL plantarflexor moment (~12%) and power (~9%) may have also contributed to the similarity in moment and positive work performed after fatigue between legs. The retention of ankle moment and work production during fatigue might speculatively be explained by the fatigue resistance of soleus, the largest muscle (by physiological cross-section) in the leg, due to its predominately slow, fatigue-resistant fibre type (Garland & McComas, 1990; Oza et al., 2017). This would have subsequently allowed energy storage in, and reuse from, the Achilles tendon when other muscle-tendon units spanning hip and knee joints reduced capacity as they fatigued. During running at a constant middle-distance pace, fatigue is primarily observed in the plantar flexors with a compensatory increase in positive work done at the knee, showing that relative joint work shifts to more proximal joints in fatigued endurance running (37, 54); these findings do not appear to extrapolate to fatigued sprinting. As such, unlike during prolonged submaximal running, there is a tendency to maintain positive and negative joint work at the ankle relative to the hip and knee during sprint running, so fatigue does not result in a more knee- or hip-dominant strategy.

Since comparatively little change was observed in DL compared to NDL after fatiguing running, the resulting similarity between limbs in fatigue can be attributed largely to a shift in kinematic and kinetic patterns in NDL. Fatigue and asymmetry are often (55–57), but not always (30, 44), cited as important yet interrelated risk factors for injury. If this is true, then the greater asymmetry observed in non-fatigued sprinting might be explained by running speed being prioritised in non-fatigued sprinting but injury risk reduction being prioritised during fatigued sprinting. Although injuries are problematic for modern athletes, injury in early humans would have had severe consequences, so the adoption of injury reduction practices would have been important. This hypothesis is worthy of explicit scrutiny in future studies.

Hamstring muscle injuries are highly prevalent in modern sports and are the most prevalent injury in running-based sports such as Association Football (soccer) (58, 59). Increases in anterior pelvic tilt during fatigued running are speculated to increase hamstrings injury risk, given that the hamstrings act across both the hip and knee joints, an increase in pelvic tilt should therefore increase hamstring muscle length (especially in the moment just prior to foot-strike; 43, 60–62). An increase in tilt of ~3° was detected in the present cohort, corroborating the findings of others (43, 60, 63). To determine whether this might have increased peak hamstrings muscle length, biceps femoris long head (BFlh), semimembranosus (SM), and semitendinosus (ST) MTU lengths were modelled in both DL and NDL (Opensim, Simtk.org, Stanford USA; see section 16 in Supplementary Information). However, no statistical differences in peak MTU lengths or the timing of peak MTU lengths were detected in any muscle in non-fatigued or fatigued sprinting, although tendencies towards small decreases in MTU length in DL but increases in NDL were observed in BFlh (~1.3%) (see Figure S14). As BFlh is the most frequently injured hamstring muscle during sprinting, and NDL (usually the kicking leg) is the most commonly injured leg (Lord et al., 2019), it might still be relevant to assess relationships between fatigue-induced BFlh length changes and injury rates in future studies.

An alternative possibility is that increased pelvic tilt provides a running speed advantage during fatigue. Anterior tilt should reduce the vertical height of the foot at the beginning of protraction, given that hip-pelvis angles (and thus muscle lengths) remain unchanged (both findings were observed in the present study). This would reduce the travel distance of the foot to the ground (also observed presently), allowing it to contact the ground before excessive fall of the CoM despite the fatigue-induced reduction in foot speed (Figure 2). The foot can therefore make contact further underneath the body rather than in front, minimising braking forces and alleviating running speed reduction. Based on this hypothesis, which requires explicit examination, anterior tilt may be a useful mechanism for maintaining running speed in the face of a decreasing foot speed.

In summary, movement pattern differences between DL and NDL indicate that the legs played partly unique roles (DL contributed more to forwards running speed) during non-fatigued sprint running in athletes who commonly perform both endurance- and sprint-type running but have not been coached in their techniques. However, differences were substantially reduced when fatigued, largely because of changes in NDL. We speculate that the increasing similarity (reduced asymmetry) in function may help to reduce injury risk in the face of fatigue, although it may also incur a cost to running speed. Additionally, rather than a shift from distal towards proximal joint work (and joint moment) being observed, as shown previously for slower running speeds, the ankle joint tended to retain its capacity for work, moment, and power production. Thus, running at faster speeds, contrary to slower-speed running, does not seem to be compromised by a loss of ankle work or power whilst in a fatigued state. Nonetheless, some limitations and delimitations are worth mentioning. Importantly, while the fatiguing running protocol was performed on a grassed sport field, sprint running testing was completed in an indoor laboratory on an athletic track (Mondo surface). While this minimised environmental effects, and the runners were given a brief re-introduction to the running track surface before post-running testing, it is also of future interest to describe sprint running mechanics on grassed and dirt surfaces, similar to those most likely used in hunter-gatherer societies and more commonly used in running-based modern sports. In addition, future work may utilise a fatiguing running protocol lasting several hours and performed on uneven terrain, more akin to that most probably performed by hunter-gatherers (but less relevant to sports) to determine whether similar results are obtained.

## Notes

### Competing Interest Statement

The authors have declared no competing interest.

