## Supplementary Information for "Fatigue does not increase limb asymmetry or induce proximal joint power shift during sprinting in habitual, multi-speed runners"

**This PDF file includes:**

1. Population and training history
    a. Rationale for cohort selection
2. Custom cluster-based marker set
3. Laboratory set-up for data collection
4. Warm-up protocol
5. Single-leg vertical jump test
6. Fatiguing running-based exercise protocol
7. Horizontal and vertical centre of mass velocity before and after fatigue
8. Ground contact time before and after fatigue
9. Discrete joint kinematics variables before and after fatigue
10. Joint work before and after fatigue
11. Joint kinematics and kinetics (statistical parametric mapping)
12. Vertical centre of mass displacement before and after fatigue
13. Discrete joint kinetics variables before and after fatigue
14. Hip extension velocity before and after fatigue
15. Visual representation of all observed statistical differences for:
    a. Non-fatigued sprinting – dominant leg and non-dominant leg
    b. Dominant leg – non-fatigued and fatigued sprinting
    c. Non-dominant leg – non-fatigued and fatigued sprinting
    d. Fatigued sprinting – dominant leg and non-dominant leg
16. Peak muscle-tendon unit length (hamstrings) before and after fatigue

### 1. Population and training history:

Thirteen semi-professional male Association Football (soccer) players (age: 19.1 ± 2.1 y, body mass: 72.5 ± 6.9 kg, height: 175 ± 7.7 cm) participated in the study. As part of their normal training and match play, the athletes regularly performed maximal speed sprinting as well as endurance-based (i.e. slow and moderate speed) running. The cohort did not receive any formal running technique instruction during training or match play. As such, the cohort represents a group of individuals who train using both lower and higher speed running on a regular basis but adopt their own implicitly learned running technique.

No subjects participated in formal strength training or other supplementary training (e.g. plyometrics, etc.) that might influence their running performance, technique, or response to fatiguing running exercise. All participants were free from injury for at least 6 months before testing and reported no residual impediments from previous injury. Only outfield players were accepted into the study as goalkeepers tend not to (or rarely) perform maximal sprinting during training or match play.

#### a. Rationale for cohort selection:

During initial study planning we had considered enrolling 800 – 3000 m track athletes into the study as they would normally perform a significant total distance of endurance running per week with smaller bouts of sprint running; they might therefore reflect the running capacities of some hunter-gatherer societies. However, all runners that we first identified had received extensive coaching in running technique (and had sought further information through less formal channels) and were specifically coached to reflect on their running technique whilst fatigued. Additionally, almost all also performed supplementary strength and/or plyometrics training, which would also have influenced their running technique and response to fatigue. Finally, it is very likely that these athletes perform far more running than in most hunter-gather societies.

We therefore broadened our view of "runners" to include those from other sports. Only field-based team sports were considered because the size of playing area in court-based sports is too small to permit athletes to (a) run for long distances uninterrupted (they rarely did this in training or games) or to reach maximal sprinting speed. We further reduced the participant scope to remove those who participated in collision sports because running technique is often changed due to accommodate impending tackles, and a lot of technique work is done during stop-start running drills (most of these athletes also perform supplementary strength and other training).

We ultimately found two accessible cohorts from which we could draw athletes who fitted our criteria: Association Football and Australian Rules football, although the latter players also participate in some collisions within the sport and tended to perform supplementary strength training. Finally, Association Football players were chosen because we were able to locate a large group of players who (i) had received no formal running instruction or training, (ii) did not perform (and had never performed) supplementary strength, plyometrics or other training, (iii) did not undergo regular collisions during training or matches, and importantly, (iv) performed sprint running efforts at maximal velocity during training and matches (i.e. under both non-stressful and stressful conditions). An important additional benefit of this choice is that Association Football is the largest participation sport in the world, so the results of the present experiments will be of broad contemporary interest.

### 2. Custom cluster-based marker set

A variety of marker set models are used for 3-D gait analyses. However, errors in model outputs can result from inaccurate marker placement and skin motion artefacts. Use of a cluster-based model has been found to reduce errors in model outputs by overcoming inaccuracies due to improper marker placement and skin motion artifacts (1), and was therefore chosen for the current study (Table S1).

**Table S1. Upper-body marker placements and naming conventions**

| Segment | Marker | Location |
|---|---|---|
| **Head** | RFHD | Right front head |
| | RBHD | Right back head |
| | LFHD | Left front head |
| | LBHD | Left back head |
| **Thorax** | C7 | Spinous process of 7th cervical vertebrae |
| | T10 | Spinous process of 10th thoracic vertebrae |
| | CLAV | Sternal notch |
| | STRN | Xyphoid process of the sternum |
| **Pelvis** | RASI | Right anterior superior iliac spine |
| | RPSI | Right posterior superior iliac spine |
| | LASI | Left anterior superior iliac spine |
| | LPSI | Left posterior superior iliac spine |
| **Right Shoulder** | RACR | Right acromion |
| **Left Shoulder** | LACR | Left acromion |

**Lower-body marker placements and naming conventions**

| Segment | Marker | Location |
|---|---|---|
| **Pelvis** | RASI | Right anterior superior iliac spine |
| | LASI | Left anterior superior iliac spine |
| | RPSI | Right posterior superior iliac spine |
| | LPSI | Left posterior superior iliac spine |
| **Right Thigh** | RTH1 | Right thigh cluster: superior marker 1 |
| | RTH2 | Right thigh cluster: superior marker 2 |
| | RTH3 | Right thigh cluster: inferior marker 1 |
| | RTH4 | Right thigh cluster: inferior marker 1 |
| **Left Thigh** | LTH1 | Left thigh cluster: superior marker 1 |
| | LTH2 | Left thigh cluster: superior marker 1 |
| | LTH3 | Left thigh cluster: inferior marker 1 |
| | LTH4 | Left thigh cluster: inferior marker 1 |
| **Right Tibia** | RTB1 | Right tibial cluster: superior marker 1 |
| | RTB2 | Right tibial cluster: superior marker 1 |
| | RTB3 | Right tibial cluster: inferior marker 1 |
| | RTB4 | Right tibial cluster: inferior marker 1 |
| | RMA | Right medial malleoli |
| | RLA | Right lateral malleoli |
| **Left Tibia** | LTB1 | Left tibial cluster: superior marker 1 |
| | LTB2 | Left tibial cluster: superior marker 2 |
| | LTB3 | Left tibial cluster: inferior marker 1 |
| | LTB4 | Left tibial cluster: inferior marker 1 |
| | LMA | Left medial malleoli |
| | LLA | Left lateral malleoli |
| **Right Foot** | RMT1 | Head of the 1$^{st}$ right metatarsal |
| | RMT5 | Head of the 5$^{th}$ right metatarsal |
| | RCAL | Right calcaneus |
| **Left Foot** | LMT1 | Head of the 1$^{st}$ left metatarsal |
| | LMT5 | Head of the 5$^{th}$ left metatarsal |
| | LCAL | Left calcaneus |

### 3. Laboratory set-up for data collection:

Thirteen VICON motion analysis cameras (Oxford Metrics Ltd., Oxford, UK) operating at a 250-Hz frame rate were used for motion capture. Data acquisition was synchronised with the ground reaction force data recordings provided by five 600 x 900-mm triaxial force platforms (Kistler Quattro, Type9290AD, Victoria, Australia) at an analogue-digital conversation rate of 1000 Hz. Motion capture cameras were positioned to create a capture volume around the in-ground force platforms over which the subjects completed their sprint running trials. This distance was chosen as team-sport athletes typically reach their maximum running speed between 30-40 m; therefore, subjects would have been at (or very near) maximum speed as they traversed the capture area (Figure S1).

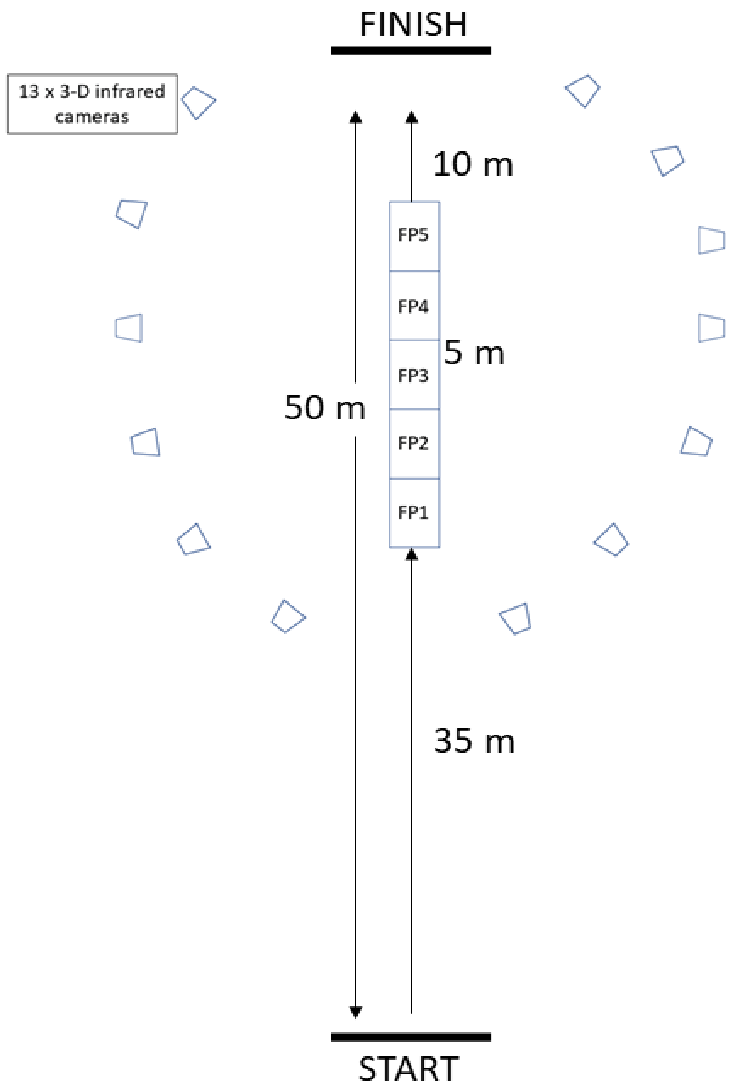


**Figure S1.** Aerial schematic of the laboratory set up during data collection. The start point was situated 35 m from the first of five serially arranged in-ground force platforms (FP 1-5). Thirteen infrared motion capture cameras were positioned around the platforms. The finishing point was designated as 10 m after the end of the capture volume to ensure subjects did not slow down within the capture area.

### 4. Warm-up protocol

Table S2 shows the progression of the standardised, comprehensive warm-up protocol completed by each participant before sprint running testing. Once the "exercise or drill" component of the warm-up was complete, the participants completed two 30-m efforts at 50% of their perceived maximal sprinting effort, one at 75% of maximal effort, and a final at absolute maximal effort. A 60-s active recovery (i.e. slow walk) was given after each effort. Table S2 shows the phase of the warm-up, exercise or drill, and prescription. A more detailed description is provided below the table for each component within the warm-up.

**Table S2.** Standardised warm protocol

| Phase | Exercise or drill | Prescription |
|---|---|---|
| **General warm-up** | Light jogging - forward, backwards, side shuffles | 5 minutes |
| | Lateral shuffle | 2 x 10 m |
| | Floor glute bridges | 1 x 8 |
| | Bodyweight squats | 1 x 10 |
| | Multi-planar lunge circuit | 1 x 6 each side |
| | Single leg squat | 1 x 5 each side |
| **Bilateral and unilateral jumps** | Single-leg hop-and-stick | 1 x 2 each side |
| | Triple hop | 1 x 2 each side |
| | Crossover hop | 1 x 2 each side |
| | Single-leg countermovement jump | 1 x 2 each side |
| **Practice runs/maximal effort** | Submaximal sprint at 50% of perceived maximum sprint effort | 2 x 30 m |
| | Submaximal sprint at 75% of perceived maximum sprint effort | 1 x 30 m |
| | Maximal sprint at 100% of perceived maximum sprint effort | 1 x 30 m |

**Exercise or drill description:**

**Lateral shuffle:** Subjects were instructed to stand with their feet hip width apart and at a self-selected pace, hinge/bend forward (i.e. flex at the hip) at the hips, bend knees (i.e. flex at the knee), looking forward, chest lifted, and neutral spine and move to the right by abducting the right leg, and as the foot reaches the ground, abduct the left leg so that the left foot finishes next to the right foot. The shuffle steps were performed with the trunk faced forward and 'hands on hips' (left and right iliac crest). This was replicated with the left side leading.

**Floor "glute bridge":** Subjects were instructed to lie down in a supine position with knees bent and feet flat on the ground with arms placed at their side on the ground. Then, to lift the buttocks off the ground until shoulders, hips, and knees formed a relatively straight line, before returning to the ground.

**Bodyweight squats:** Subjects were instructed to stand with their feet hip width apart and to squat down to a self-selected depth while looking forward, chest lifted, and neutral spine. Then, extend the hip and knee to return to standing position. This was performed at a self-selected pace.

**Multi-planar lunge circuit:** Subjects were instructed to perform lunges in the sagittal, frontal, and transverse planes. All three lunges started with the subjects standing with their feet slightly closer than hip width apart. Each lunge was performed keeping the trunk in an upright position so that the knee and hip of the leading leg flexed to ~90°.

- Sagittal plane lunge: step forward with one leg, once the foot reaches the ground, lower the body toward the ground, then push off the front foot and return to starting position.
- Lateral lunge: step laterally with one foot, once the foot reaches the ground ensure majority of the bodyweight is supported with that leg, flex at the hip and push buttocks backwards while the stationary leg extends at the knee. Then push off the lateral leg and return to starting position.
- Transverse lunge: Subjects were instructed to step laterally and rotate (trunk and leg) away from the stationary leg; the movement was instructed to be performed with rotation and lunge in a consecutive motion. Then push off the lateral leg and return to starting position.

### 5. Single-leg vertical jump:

Sprinting at maximum velocity necessitates rapid development of forces in the lower limbs whilst moving at high speeds (6–8). Therefore, a performance test that specifically examines force production whilst trying to move rapidly should best define legs according to 'strongest' and 'weakest', or in this case, dominant and non-dominant. In some studies, researchers have designated the preferred kicking leg as dominant, which may not be the stronger of the two legs and thus may not accord with our definition.

Unlike in the upper limbs, limb dominance in the lower limbs is defined inconsistently between studies and varies as the leg chosen for kicking, strength, perception, or task-specific capacity (9, 10). Whilst the kicking leg is often defined as the dominant limb, we chose to measure lower limb force production capacity rather than to use the preferred kicking leg because the leg chosen for kicking is rarely the strongest leg or used as the take-off leg when jumping in a match, and therefore is unlikely to produce the most force rapidly during sprinting (11). Movements requiring significant skill, such as single-leg jumping or hopping, tend to have larger interlimb differences and are therefore less ambiguous in identifying the dominant and non-dominant limbs (12). As such, single-leg vertical jumps were chosen to identify the dominant and non-dominant limbs.

Typically, higher intensities and longer durations of exercise cause greater levels of fatigue and consequently induce greater impairment in muscle function compared to less intense or lower volume exercises (13, 14). Fatigue-induced impairment tends to result in decreased force production capability and jump height in vertical jump tests, which has been used as a reliable measure to detect acute fatigue (15, 16). The single-leg vertical jump test is a suitable tool to detect fatigue-induced changes, with several variables associated with very high reproducibility (Edwards et al., 2018; Gathercole et al., 2015).

To evaluate force production capability of each limb as a method of determining limb dominance and the detection of fatigue-induced changes, net impulse was calculated by numerically integrating the vertical force using the trapezoid rule (19). Then, vertical net impulse was divided by body mass to determine the take-off velocity, and jump height was calculated from the take-off velocity by the following equation, where $TOV^2$ represents the square of take-off velocity, $g$ represents the acceleration of gravity (9.81 m·s$^{-2}$): $Jump\ height = \frac{TOV^2}{2g}$. The limitation of using impulse in isolation is that it can be increased by either applying a larger (peak) force or by extending the time over which force is produced, though long force production durations are associated with reduced jump height (15). Therefore, a combination of vertical net impulse and vertical jump height has been used to provide a reliable and valid measure of force production capability for each limb (20).

Subjects performed three single-leg vertical jumps each with hands on placed hips to minimise the influence of arm swing. Figure S2 shows the mean single-leg vertical jump heights and Table S3 shows the net vertical impulse for each subject for the dominant and non-dominant legs in the non-fatigued and fatigued trials. Paired t-tests identified significant differences in jump height ($p < 0.001$) and net vertical impulse ($p < 0.013$) between DL and NDL in the non-fatigued condition, with DL ~1.6 cm greater jump height and ~8% greater vertical net impulse than NDL. After fatiguing exercise, no statistical differences were observed between DL and NDL ($p < 0.069$; $p < 0.111$). Significant differences in jump height were observed between non-fatigued and fatigued conditions for DL ($p < 0.001$) and NDL ($p < 0.001$), and no differences in net vertical impulse (DL - $p < 0.081$; NDL - $p < 0.431$). The calculation of jump height and vertical net impulse during the single-leg vertical jump test clearly identified the limb that produced greater force, which was therefore defined as the dominant limb. After fatiguing exercise, both limbs decreased in the selected variables, indicating sufficient sensitivity to detect fatigue-induced changes in the single-leg muscle function. This confirms that, after completion, the Ball-Sport and Endurance Test protocol induced detectible fatigue within the cohort.

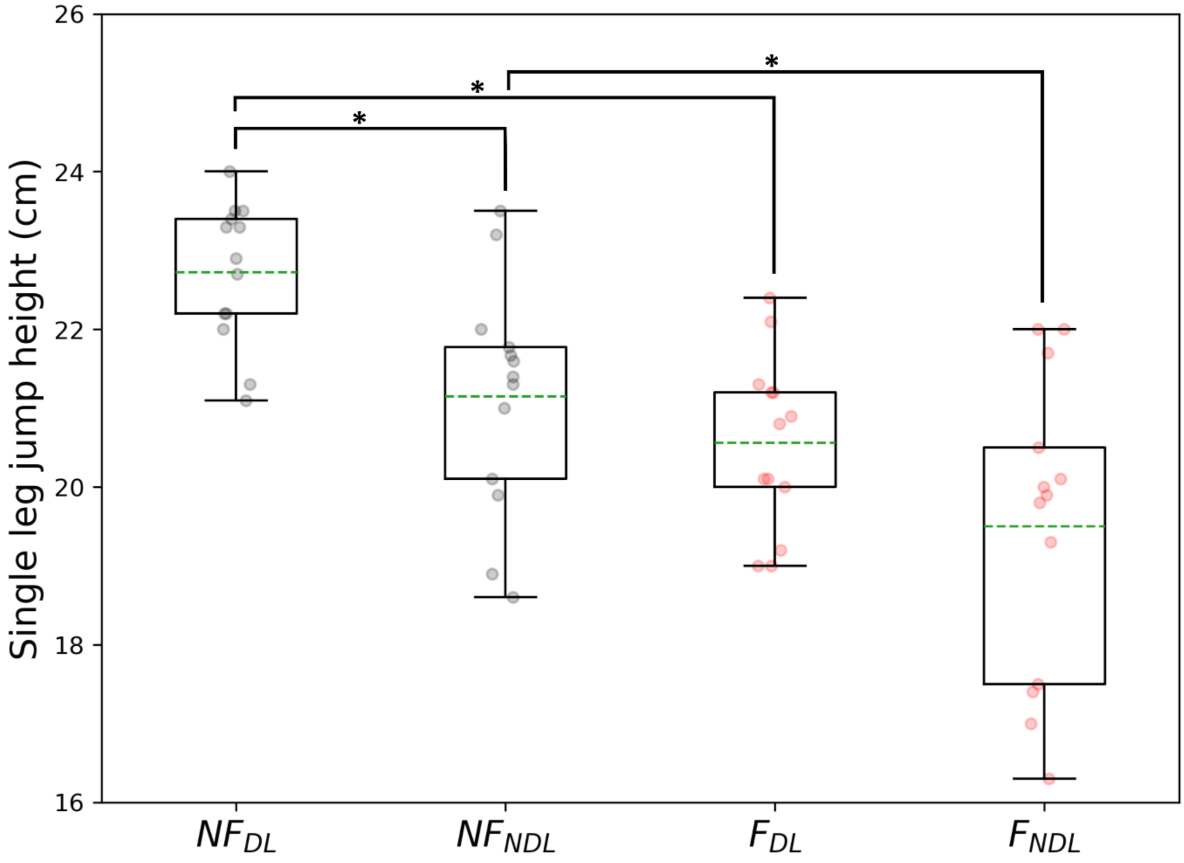


**Figure S2.** Mean single-leg vertical jump heights are shown for the non-fatigued (NF; grey circles) and fatigued (F; red circles) conditions in the dominant (DL) and non-dominant (NDL) legs. *statistical difference between legs or conditions ($p < 0.05$). Green dashed line = mean value for all participants.

**Table S3.** Non-fatigued and fatigued vertical impulse for the dominant and non-dominant legs. * statistical difference between the dominant and non-dominant legs trials. ^ statistical difference between the non-fatigued and fatigued trials, respectively.

| | **Non-fatigued** | **Fatigued** | **95% CI** |
|---|---|---|---|
| | **Mean ± SD** | **Mean ± SD** | |
| ***Dominant leg*** | | | |
| Vertical impulse (kg·m/s) | 0.79 ± 0.11*^ | 0.74 ± 0.11* | 0.74, 0.85 |
| ***Non-dominant leg*** | | | |
| Vertical impulse (kg·m/s) | 0.74 ± 0.13^ | 0.72 ± 0.09 | 0.67, 0.77 |

### 6. Fatiguing running-based exercise protocol – Ball-Sport Endurance and Sprint Test (BEAST - 45 min)

After performing the pre-fatigue sprints, subjects completed the protocol shown below. Subjects jogged for 100 m at a comfortable pace to a level, grassed sports surface (football pitch) where markers demarcated the BEAST protocol. The beast protocol has been shown to be a valid and reliable simulator of soccer match play with respect to movement time, movement patterns, and physical demands. In addition, there is evidence that this protocol induces similar levels of fatigue to that found during a match (Delextrat et al., 2018; Matthews et al., 2017; Williams et al., 2014; Williams et al., 2010). The testing protocol involved sprinting, jogging, walking, jumping, and backwards running. Subjects were verbally encouraged to adhere to all instructions, i.e. to run as fast as possible during the sprint sections and to decelerate within the allocated areas. This protocol was familiar to the participants, who were practiced in completing it without a notable pacing strategy and did not require further, extensive familiarisation. The protocol requires all directions of movement that might be needed in the hunt or chase of an agile animal and is also reflective of the movement patterns required in many modern sports.

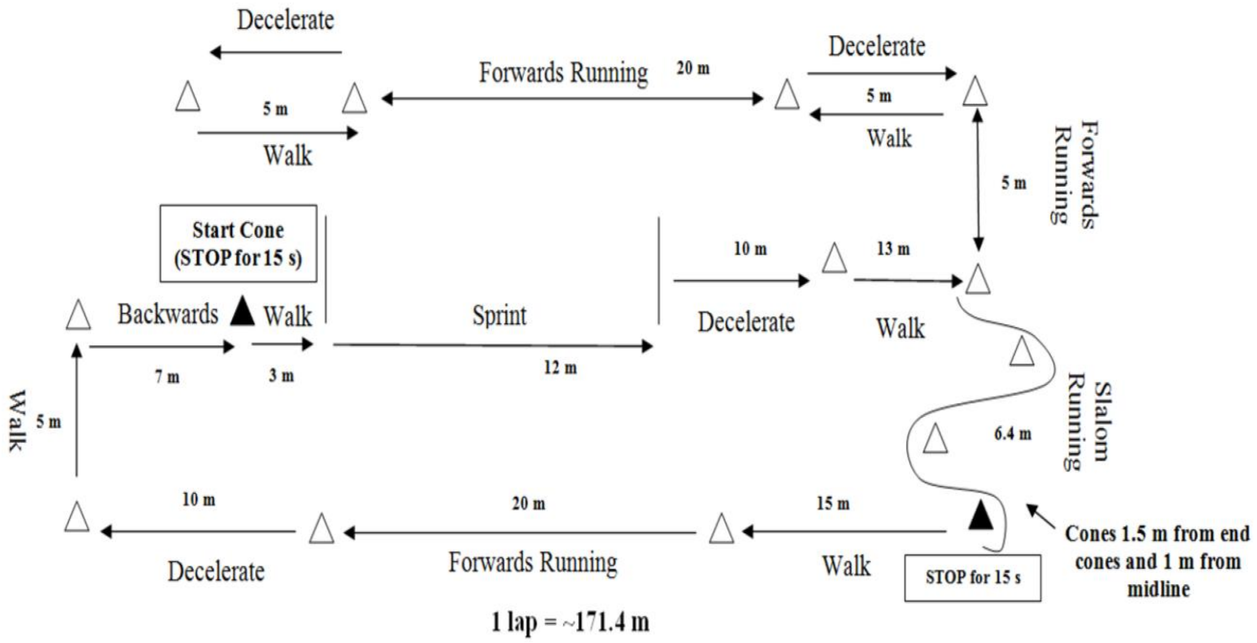


**Figure S3.** Schematic of the Ball-sport Endurance Test (BEAST) – 45 min.

### 7. Centre of mass (whole-body) velocity

The horizontal and vertical centre of mass (CoM) velocities tended to vary slightly throughout the sprinting cycle. In the second half of the stance phase, the CoM is projected forward and upward into the flight phase allowing the lower limbs to reposition for the subsequent foot-strike (Mann & Hagy, 1980; von Lieres und Wilkau et al., 2020). Large stride-to-stride differences in horizontal and vertical CoM velocity reduces running efficiency (Mann & Sprague, 1980; Slawinski et al., 2010), for example, if the vertical CoM slows down to a greater extent, then it will subsequently require greater re-acceleration to maintain running velocity.

CoM velocity was obtained from the scaled skeletal model in Visual 3D (C-Motion, Germantown, MD, USA). Table S4 shows the maxima, minima, mean difference, and 95% confidence intervals of the changes in centre of mass (CoM) velocity during sprinting trials in the dominant (DL) and non-dominant (NDL) legs in non-fatigued and fatigued conditions. Paired t-tests identified significant differences between non-fatigued and fatigued sprinting for both DL and NDL as maximum horizontal CoM velocity decreased (-0.3 m/s). Additionally, differences were observed in minimum horizontal CoM velocity between legs in the fatigued condition. That is, the horizontal CoM velocity decreased more in NDL than DL with fatigue. The minimum vertical CoM velocity decreased for NDL when fatigued; conversely, no changes were observed for DL. *statistical difference between non-fatigued and fatigued trials. ^statistical difference between DL and NDL.

**Table S4.** Maxima, minima, mean difference, and 95% confidence interval of the change in centre of mass (CoM) velocity in non-fatigued and fatigued conditions in the dominant leg.

| | Non-fatigued | Fatigued | Mean diff | 95% CI (change) |
|---|---|---|---|---|
| | Mean ± SD | Mean ± SD | | |
| ***Dominant leg*** | | | | |
| Max horizontal $V_{CoM}$ | 8.56 ± 0.47* | 8.21 ± 0.48* | -0.34 | 0.02, 0.66 |
| Min horizontal $V_{CoM}$ | 7.57 ± 0.63* | 7.21 ± 0.45*^ | -0.35 | 0.03, 0.67 |
| Mean difference (max-min) | 0.99 ± 0.25 | 1.00 ± 0.29 | | |
| Max vertical $V_{CoM}$ | 0.39 ± 0.15 | 0.42 ± 0.16 | -0.03 | -0.12, 0.07 |
| Min vertical $V_{CoM}$ | -0.52 ± 0.15 | -0.56 ± 0.15 | -0.01 | -0.79, 0.07 |
| Mean difference (max-min) | 0.91 ± 0.27 | 0.96 ± 0.21 | | |
| ***Non-dominant leg*** | | | | |
| Max horizontal $V_{CoM}$ | 8.62 ± 0.42* | 8.29 ± 0.42* | -0.33 | 0.14, 0.52 |
| Min horizontal $V_{CoM}$ | 7.49 ± 0.56* | 6.95 ± 0.52*^ | -0.54 | 0.30, 0.79 |
| Mean difference (max-min) | 1.13 ± 0.37 | 1.35 ± 0.48 | | |
| Max vertical $V_{CoM}$ | 0.42 ± 0.17 | 0.46 ± 0.16 | -0.04 | -0.12, 0.05 |
| Min vertical $V_{CoM}$ | -0.48 ± 0.19* | -0.56 ± 0.15* | 0.09 | 0.02, 0.15 |
| Mean difference (max-min) | 0.9 ± 0.26 | 1.02 ± 0.21 | | |

### 8. Ground contact times

Ground contact times in elite sprinters are typically <0.1 s when running in track spikes on a hard, elastic rubber-based track (Mann & Hagy, 1980; Morin et al., 2015). In our cohort, contact times ranged 0.110 – 0.155 s across both conditions (Mann & Hagy, 1980; Morin et al., 2015). Shorter ground contact times are commonly associated with faster maximal sprinting velocity, indicating that elite sprinters are able to generate large ground forces rapidly (26), so it was not unexpected that contact times would be slightly longer for the current cohort of untrained (although experienced) runners. Ground contact times were similar between legs in the non-fatigued condition (mean DL = 0.133 s; NDL = 0.137 s). Times statistically increased in the fatigued condition ($p < 0.012$) in NDL only. Additionally, comparison between legs showed that ground contact times increased ~4% more in NDL than DL from non-fatigued to fatigued conditions.

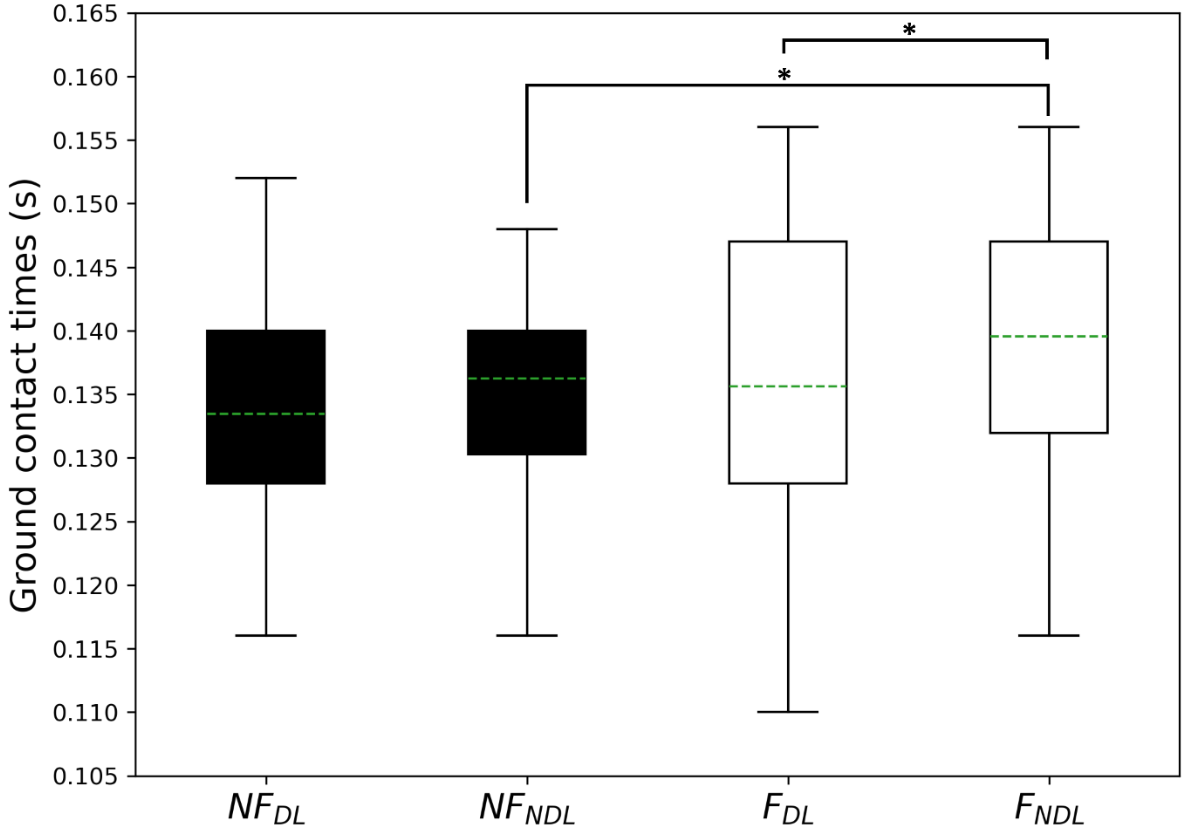


**Figure S4.** Boxplots representing foot-ground contact times in non-fatigued (NF) and fatigued (F) conditions for dominant (DL) and non-dominant (NDL) legs. *statistical difference between legs or conditions. Dashed green line = mean value of the cohort.

### 9. Peak joint kinematics

**Table S5.** Discrete kinematic variables for the pelvis, hip, and knee joints during the retraction and protraction phases for the non-fatigued and fatigued in dominant and non-dominant legs. *statistical difference between non-fatigued and fatigued trials for dominant and non-dominant legs. ^statistical difference between in the changes in dominant and non-dominant legs for non-fatigued and fatigued trials, respectively.

| | Non-fatigued | Fatigued | Mean diff | 95% CI (change) |
|---|---|---|---|---|
| | Mean ± SD | Mean ± SD | | |
| ***Dominant leg*** | | | | |
| **Pelvis angle (°)** | | | | |
| Peak anterior tilt during early protraction | 0.1 ± 8.6* | 3.6 ± 8.4* | 3.6 | 1.0, 7.0 |
| Peak anterior tilt at foot-strike | 2.5 ± 9.7 | 3.0 ± 7.8 | 0.4 | -1.5, 2.4 |
| **Hip angle (°)** | | | | |
| Peak flexion angle during retraction | 65.8 ± 6.0^ | 65.2 ± 7.2 | 0.6 | -3.9, 5.2 |
| Angle at foot-strike | 41.6 ± 7.3 | 42.6 ± 7.4 | -1.0 | -2.7, 0.6 |
| Angle at toe-off | -7.4 ± 8.2 | -6.4 ± 10.5 | -1.0 | -3.9, 2 |
| **Knee angle (°)** | | | | |
| Peak extension angle prior to foot-strike | -45.8 ± 11 | -45.7 ± 14.4 | -0.1 | -10.8, 10.6 |
| Angle at foot-strike | -25.2 ± 5.4 | -24.3 ± 5.3 | -0.9 | -3.4, 1.7 |
| Angle at toe-off | -20.1 ± 4.3 | -21.7 ± 6.9* | 1.5 | -3.4, 6.4 |
| ***Non-dominant leg*** | | | | |
| **Pelvis angle (°)** | | | | |
| Peak anterior tilt during early protraction | 0.8 ± 9.7* | 3.2 ± 9.9* | 2.5 | 1.0, 4.0 |
| Peak anterior tilt at foot-strike | 1.8 ± 7.4 | 1.6 ± 7.0 | -0.2 | -1.2, 0.8 |
| **Hip angle (°)** | | | | |
| Peak flexion angle during retraction | 72.2 ± 9.0*^ | 69.0 ± 10.0* | 3.0 | 1.2, 4.7 |
| Angle at foot-strike | 45.0 ± 6.8 | 43.7 ± 7.4 | 1.3 | -3.9, 6.4 |
| Angle at toe-off | -10.7 ± 3.8* | -12.8 ± 3.8* | 2.1 | 0.0, 4.2 |
| **Knee angle (°)** | | | | |
| Peak extension angle prior to foot-strike | -54.0 ± 16.6* | -46.4 ± 15.2* | -7.6 | -11.4, -3.8 |
| Angle at foot-strike | -25.6 ± 6.1 | 23.7 ± 6.2 | -1.9 | -5.3, 1.5 |
| Angle at toe-off | -17.2 ±3.1 | -15.8 ± 3.2* | -1.4 | -3.6. 0.7 |

### 10. Joint work

Inverse dynamic analyses were used to compute net joint moments, which were subsequently multiplied by joint angular velocities to obtain joint powers at the hip, knee, and ankle joints. To obtain positive and negative mechanical work performed by the lower limbs, joint power data were individually integrated with respect to time using the trapezoidal method. For each limb, all values of positive work were summed, and all negative work values were summed, to give individual joint totals for positive and negative work, respectively.

The average positive powers calculated for the hip, knee, and ankle joints were summed and this value was described as total positive and power output (equation 1) where $P_{tot}^{+}$, $P_{hip}^{+}$, $P_{knee}^{+}$, $P_{ankle}^{+}$ are total, hip, knee, and ankle joint average positive powers. Each joint's average positive power as a percentage of total average positive power was determined (equation 2), where $J_{percent}$ is the percentage of an individual joint to the total work. The same equations were used to obtain total average negative power for each joint.

(1) $$P_{tot}^{+} = P_{hip}^{+} + P_{knee}^{+} + P_{ankle}^{+}$$

and

(2) $$J_{percent} = \left(\frac{P_j^{+}}{P_{tot}^{+}}\right) \times 100\%$$

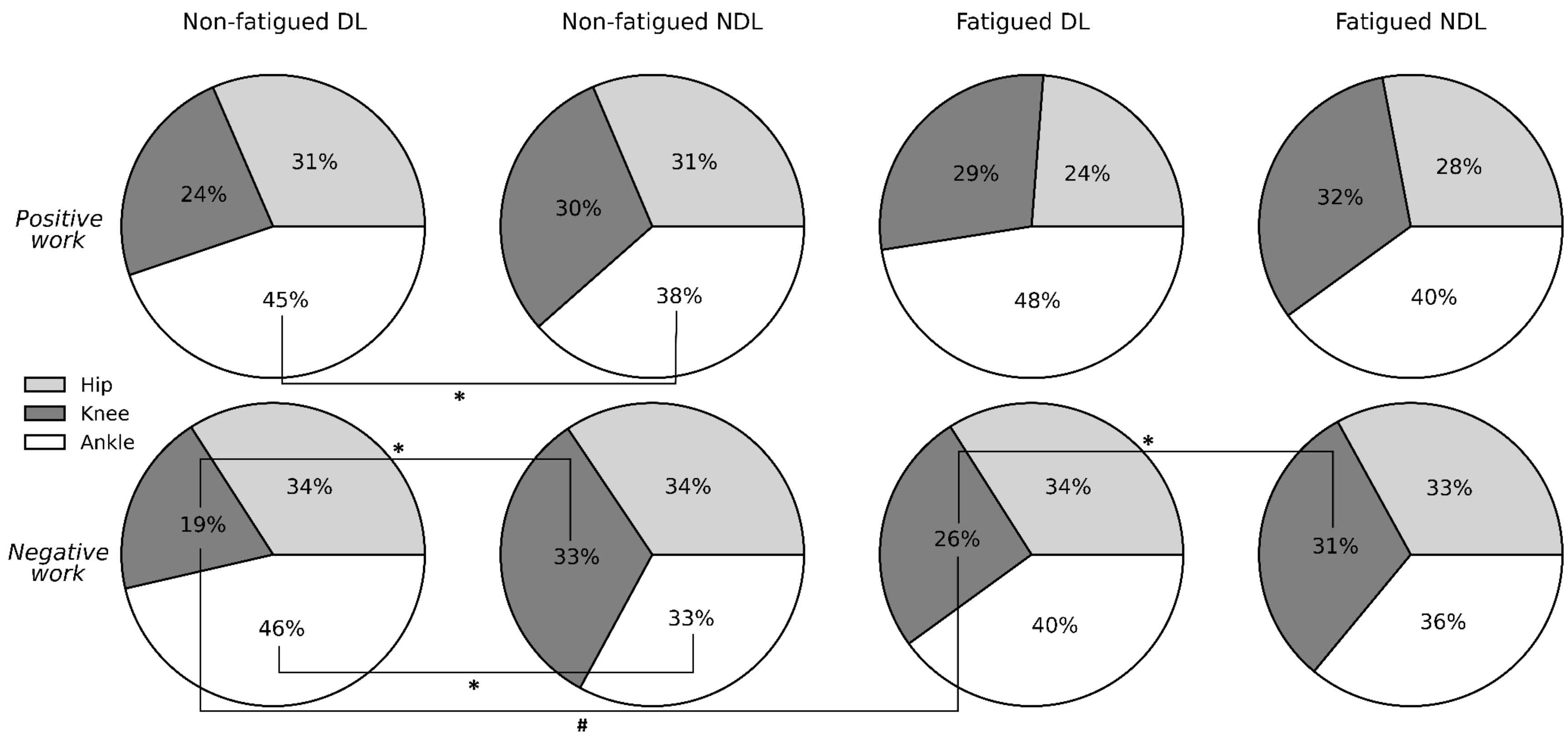


**Figure S5.** Percentage of total average positive and negative power contributed by the hip (light grey), knee (dark grey), and ankle (black) joints in dominant (DL) and non-dominant (NDL) legs. Significant differences were observed in DL and NDL positive ankle work, negative ankle work, and negative knee work in non-fatigued sprinting, but only in negative knee work in fatigued sprinting, despite statistical increase (#) in DL after fatigue. * statistical difference between legs within conditions. # statistical change between non-fatigued and fatigued conditions ($p < 0.05$).

### 11. Peak joint kinetics

**Table S6.** Kinetic variables for the hip, knee, and ankle joints of the dominant and non-dominant legs in the non-fatigued and fatigued conditions. * statistical difference between non-fatigued and fatigued trials for dominant and non-dominant legs. ^ statistical difference between dominant and non-dominant legs in the non-fatigued and fatigued conditions, respectively.

| | Non-fatigued | Fatigued | Mean diff | 95% CI (change) |
|---|---|---|---|---|
| | **Mean ± SD** | **Mean ± SD** | | |
| ***Dominant leg*** | | | | |
| **Hip** | | | | |
| Peak Extension Moment | -3.6 ± 0.7 | -3.4 ± 0.4 | -0.2 | -0.7, 0.3 |
| Peak Power (Generated) | 30.7 ± 10.2 | 27.2 ± 7.8 | 3.5 | -2.5, 9.5 |
| **Knee** | | | | |
| Peak Extension Moment | 3.3 ± 0.9*^ | 3.7 ± 1.0* | -0.4 | 0.0, 0.6 |
| Peak Power (Generated) | 14.8 ± 6.0^ | 13.2 ± 6.5 | -1.6 | -5.3, 5.9 |
| **Ankle** | | | | |
| Peak Plantarflexion Moment | -3.4 ± 0.6^ | -3.2 ± 0.8 | -0.2 | -0.7, 0.4 |
| Peak Power (Generated) | 26.4 ± 7.1 | 23.3 ± 8.7 | -3.1 | -2.1, 8.4 |
| ***Non-dominant leg*** | | | | |
| **Hip** | | | | |
| Peak Extension Moment | -3.6 ± 0.6* | -3.2 ± 0.5* | -0.4 | -0.6, 0.0 |
| Peak Power (Generated) | 32.6 ± 8.9 | 28.9 ± 6.1 | -3.7 | 0.4, 7.1 |
| **Knee** | | | | |
| Peak Extension Moment | 3.7 ± 0.7^ | 3.7 ± 0.6 | 0.1 | -0.2, 0.0 |
| Peak Power (Generated) | 18.3 ± 7.3*^ | 15.5 ± 6.8* | -2.8 | -6.3, 2.1 |
| **Ankle** | | | | |
| Peak Plantarflexion Moment | -3.0 ± 0.8^ | -3.2 ± 0.8 | 0.2 | 0.0, 0.4 |
| Peak Power (Generated) | 24.1 ± 8.0 | 25.9 ± 7.3 | -1.8 | -4.3, 0.7 |

### 12. Statistical Parametric Mapping - joint kinematics and kinetics

Statistical parametric mapping (SPM) is a topological methodology for detecting field changes that are continuous functions of space or time (27). Many classes of biomechanical data are smooth and contained within bounds, and as such are well suited to SPM analyses. We used SPM as it provides statistical significance for individual points and/or regions within the sprint cycle.

Here, we compared the hip, knee, and ankle joint kinematics and kinetics of the dominant leg (DL) during non-fatigued and fatigued sprinting (Figure S6). The SPM analyses did not detect significant statistical changes at the hip and negligible difference in the ankle joint (i.e. a collection of ≥5 consecutive points exceeding the threshold did not occur (28)) between non-fatigued and fatigued sprinting. Significant differences were observed in knee joint kinetics (moments and power) from around foot-ground contact to early stance (8 consecutive points exceeded the threshold).

The SPM analyses did not detect significant changes at the hip and knee joints of the non-dominant leg (NDL) between non-fatigued and fatigued sprinting. However, an unexpected, yet significant difference was observed in ankle joint kinetics (moments and power) from around foot-ground contact to early stance (18 consecutive points exceeded the threshold). That is, during fatigued sprinting, the ankle joint produced significantly greater ankle plantarflexion moments as well as negative joint power (Figure S7).

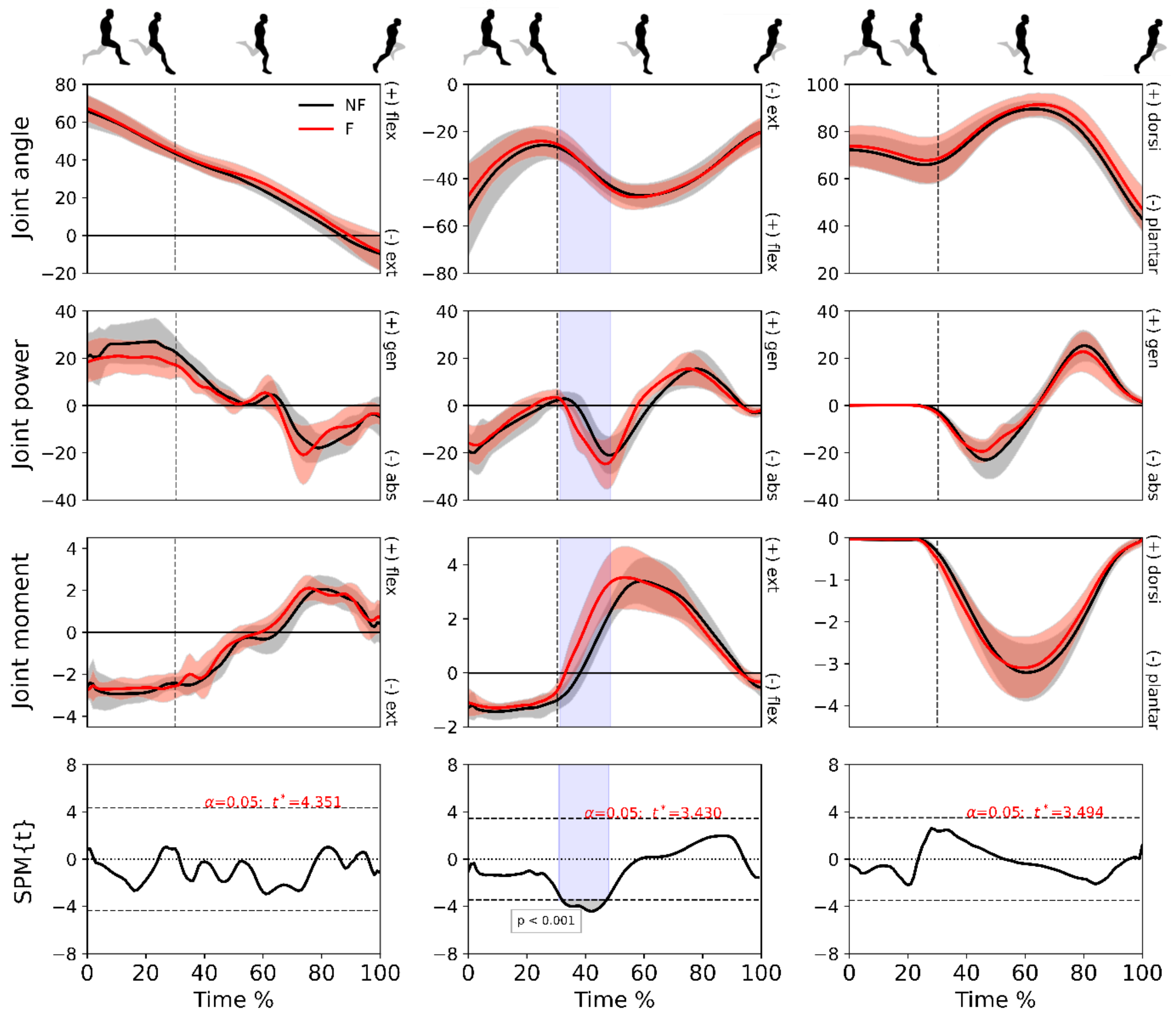


**Figure S6.** Mean (± standard deviation) joint angles (degrees), powers (W/kg) and moments (N/kg) at hip (column 1), knee (column 2) and ankle (column 3) joints for the dominant leg only in the non-fatigued (black) and fatigued (red) conditions. Vertical dotted line represents foot-strike. The time-dependent paired t-values of the SPM (bottom row; set at $p < 0.05$) are shown as horizontal dashed lines. Shaded areas indicate regions with statistical differences.

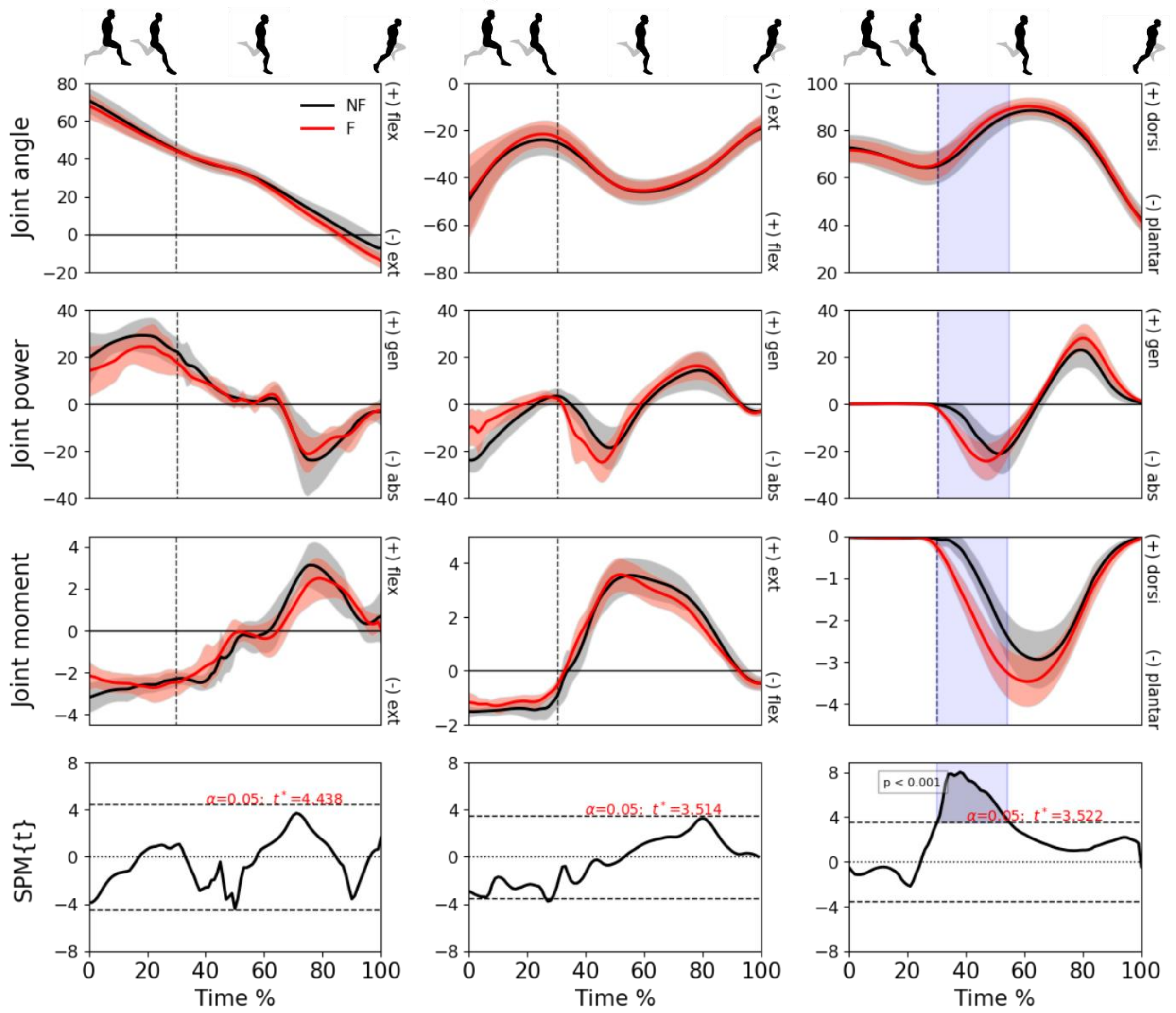


**Figure S7.** Mean (± standard deviation) joint angles (degrees), powers (W/kg) and moments (N/kg) at hip (column 1), knee (column 2) and ankle (column 3) joints for the non-dominant leg only in the non-fatigued (black) and fatigued (red) conditions. Vertical dotted line represents foot-strike. The time-dependent paired t-values of the SPM (bottom row; set at $p$ < 0.05) are shown as horizontal dashed lines. Shaded areas indicate regions with statistical differences.

### 13. Vertical centre of mass displacement

The magnitude of vertical centre of mass (CoM) displacement is strongly associated with running economy. Reducing vertical displacement improves running economy by reducing metabolic cost associated with mechanical loading of the lower limbs (29). All movements diverging from running direction negatively affect running economy, especially so in vertical displacement of the CoM (Gullstrand et al., 2009; Thorstensson et al., 1984).

A similar vertical CoM displacement was observed for DL and NDL in the non-fatigued condition, but statistical differences were observed for NDL between the non-fatigued and fatigued conditions ($p < 0.049$), and in the fatigued condition between DL and NDL ($p < 0.020$) (Figure S8).

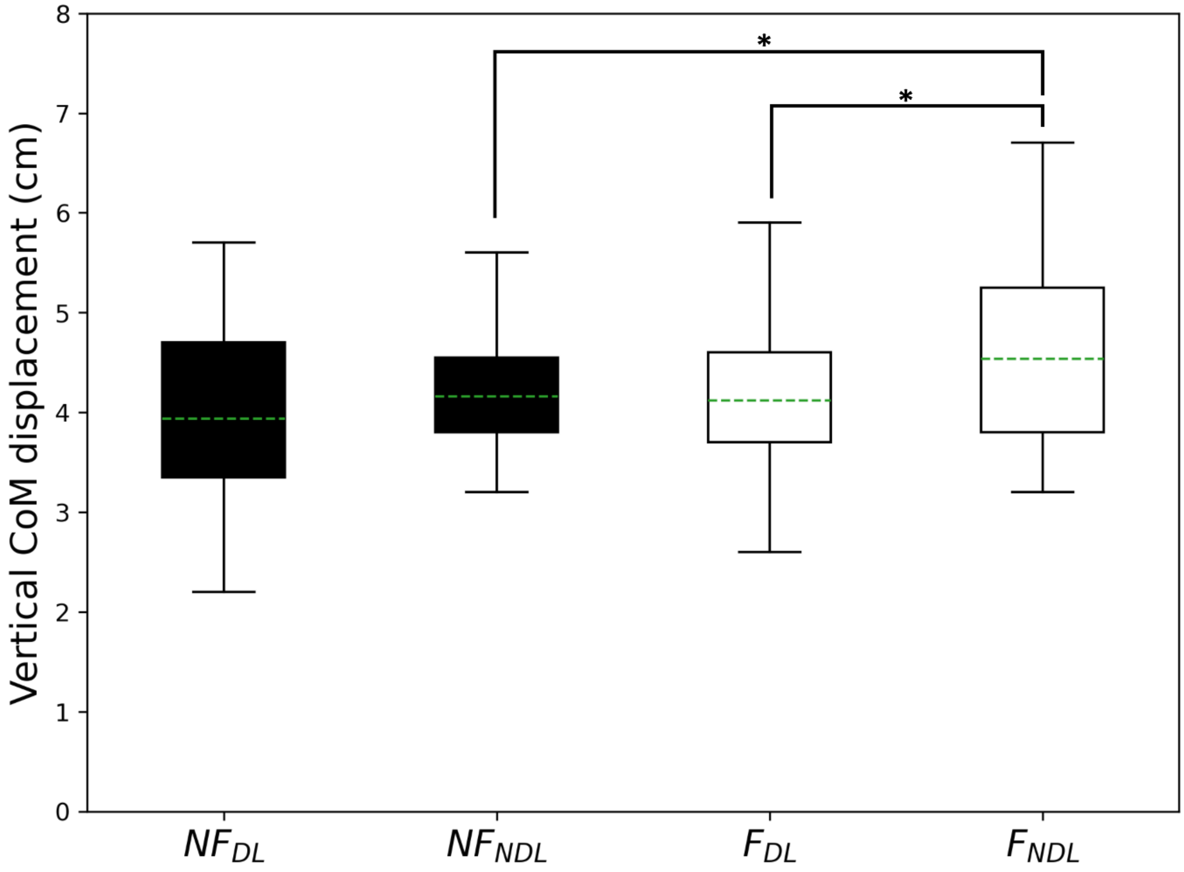


**Figure S8.** Data shown for the vertical centre of mass displacement in centimetres from the retraction and protraction phases. Data is shown for the non-fatigued (NF) and fatigued (F) conditions for the dominant (DL) and non-dominant legs (NDL). * denotes statistical significance between the non-fatigued and fatigued trials for the dominant leg. Dashed green line represents the mean.

### 14. Peak hip extension velocity

As running speed increases above ~7 m/s, increases in horizontal velocity are result from increases in stride frequency (32). The muscles surrounding the hip (and, to a lesser extent, the knee) achieve this by accelerating the foot down toward the ground more rapidly (26). Previous studies have shown greater vertical force production is normally associated with increases in hip extension angular velocity and vertical velocity of the foot, so the faster the hip extends to accelerate the lower limb down toward the ground, the more force is produced vertically. In our cohort, although peak hip extension velocity varied individually and changes with fatigue did not reach statistical significance, the larger vertical impulse produced by NDL may partly be explained by ~24% greater peak hip angular extension velocity (Table S7).

**Table S7.** Peak hip extension velocity for the dominant and non-dominant legs for the non-fatigued and fatigued. * statistical difference between non-fatigued and fatigued conditions for dominant and non-dominant legs. ^ statistical difference between dominant and non-dominant legs for non-fatigued and fatigued conditions, respectively

| | **Non-fatigued** | **Fatigued** | **Mean diff** | **95% CI (change)** |
|---|---|---|---|---|
| | **Mean ± SD** | **Mean ± SD** | | |
| ***Dominant leg*** | | | | |
| **Peak hip extension velocity (deg/s)** | | | | |
| Early protraction phase | -478 ± 131 | -468 ± 137 | -10.0 | 56, 189 |
| ***Non-dominant leg*** | | | | |
| **Peak hip extension velocity (deg/s)** | | | | |
| Early protraction phase | -627 ± 108 | -571 ± 110 | -56.0 | 88, 201 |

### 15. Visual representation of all statistical differences observed (Figures S9-12)

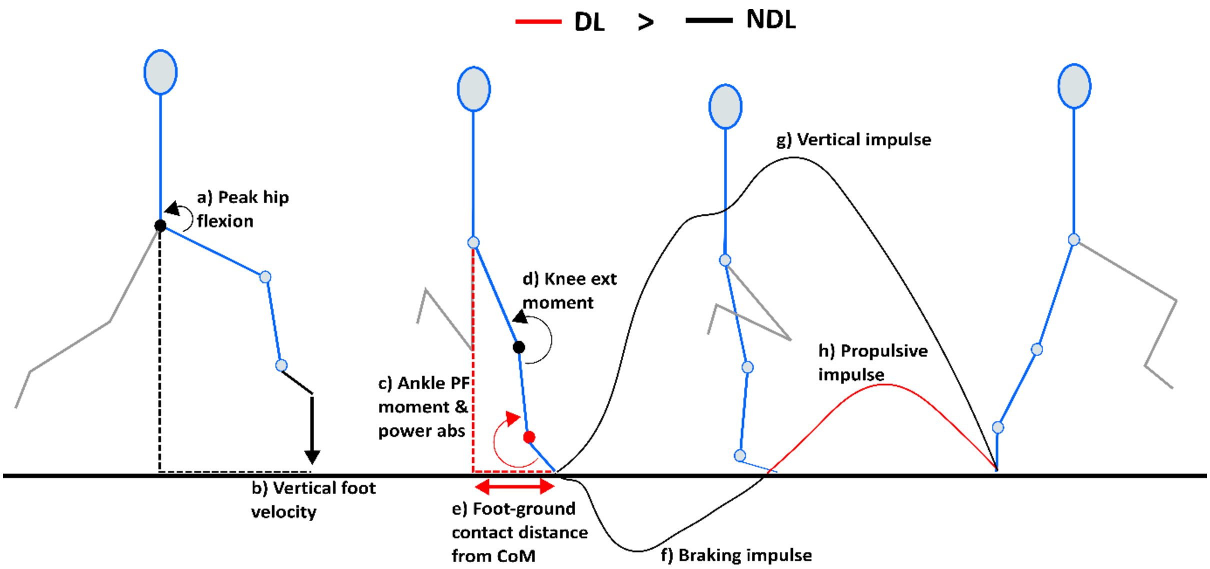


**Figure S9.** Visual representation of the differences observed between the dominant are displayed in red and non-dominant leg in black in non-fatigued sprinting. The statistical differences observed in the kinematic and kinetic variables for the dominant leg are displayed in red and non-dominant leg in black. NDL had greater peak hip flexion angle (a) and vertical velocity of the foot (b) toward the ground, larger knee extension moment (d), as well as larger braking (f) and vertical impulse (g). DL had greater ankle (kinetic) contribution (c), foot-ground contact position more underneath the centre of mass (e), and greater propulsive impulse (h).

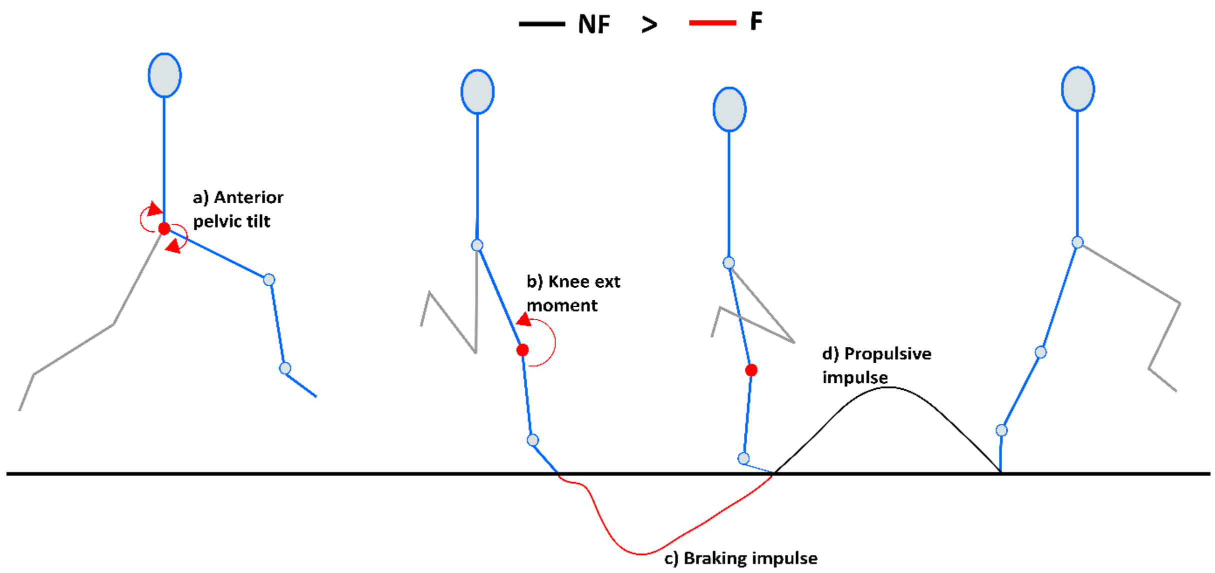


**Figure S10.** Visual representation of the differences observed the dominant leg in non-fatigued and fatigued sprinting. The statistical differences observed in the kinematic and kinetic variables for the non-fatigued condition are displayed in black and the fatigued condition in red. After fatiguing exercise, anterior pelvic tilt increased (a), greater knee extension moment (b) and braking impulse (c) were also observed. Greater propulsive impulse (d) was observed in non-fatigued sprinting;

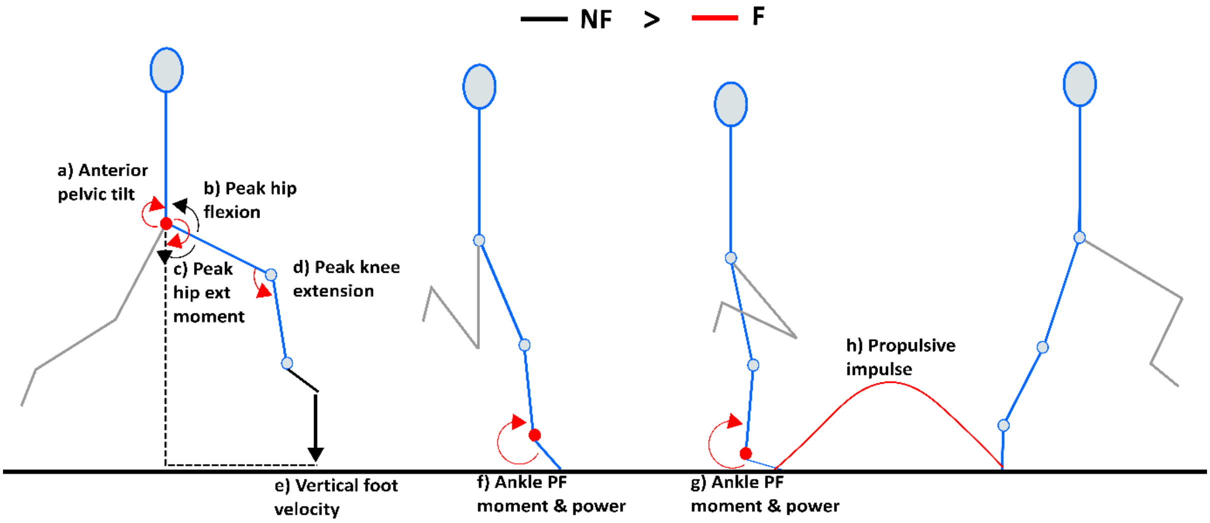


**Figure S11.** Visual representation of the differences observed the non-dominant leg in non-fatigued and fatigued sprinting. The statistical differences observed in the kinematic and kinetic variables for the non-fatigued condition are displayed in black and the fatigued condition in red. After fatiguing exercise, anterior pelvic tilt increased (a), more extended knee angle (d) in early protraction, greater ankle (kinetic) contribution (f-g), and greater propulsive impulse (h). Greater peak hip flexion (b) and vertical velocity of the foot relative to the centre of mass (e) was observed in non-fatigued sprinting.

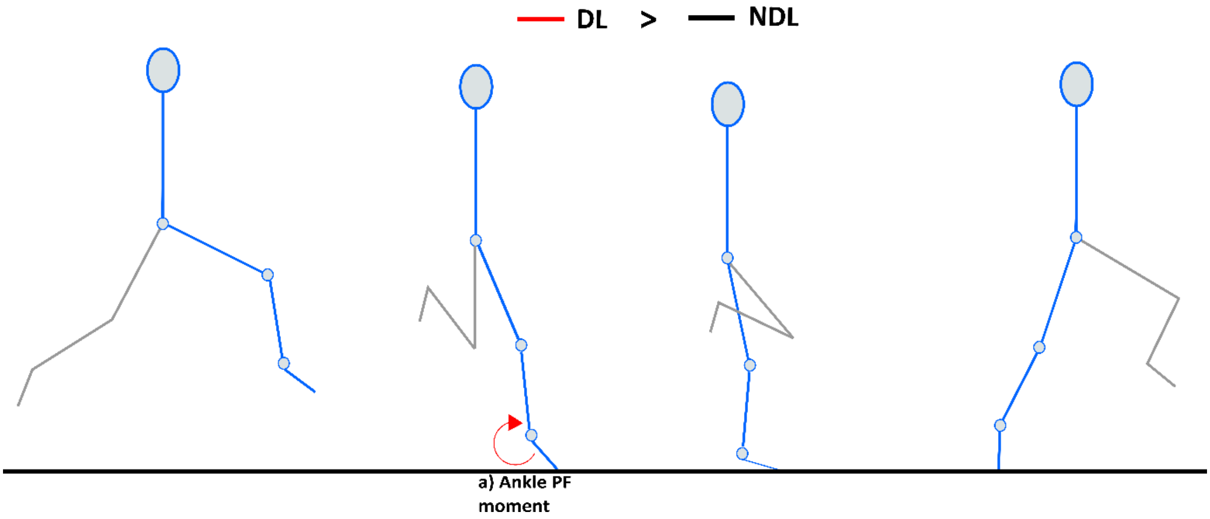


**Figure S12.** Visual representation of the differences observed between the dominant and non-dominant leg in fatigued sprinting. The statistical differences observed in the kinematic and kinetic variables for the dominant leg are displayed in red and non-dominant leg in black. After fatiguing exercise, besides a slightly greater ankle (kinetic) contribution (a), no other significant differences were observed.

### 16. Peak muscle-tendon unit length (hamstrings):

The hip transitions from flexion to extension early in the protraction phase during maximal speed sprinting. Subsequently, the knee extends as the foot travels toward the ground; here, the proximal hamstrings contract to extend the hip whilst the distal ends are subjected to strain as the knee extends prior to foot-ground contact (33–35). Although there tends to be inter-muscle differences in both stretch and lengthening velocity of the whole hamstrings muscle-tendon unit (MTU), current evidence indicates that peak MTU length may be a primary predictor of hamstring injury (36). Peak MTU length occurs just prior to foot-strike and tends to be larger in the biceps femoris long head compared to the other hamstring muscles (37). An increase in anterior pelvic tilt (APT) during sprinting has been speculatively linked to hamstring injury (38). Hypothetically, if the hip and knee angles remain similar while anterior pelvis tilt increases then the working length of the hamstring muscle-tendon unit should increase, thus increasing strain injury risk. The present results revealed a greater anterior pelvic tilt just prior to foot-strike for both DL (~3.6°) and NDL (~2.5°) when fatigued, so we were able to test this hypothesis.

The Visual 3D (C-Motion, Germantown, MD, USA) software program allows the user to export OpenSim-compatible motion files designed for use with OpenSim gait models (Simtk.org, Stanford USA). Opensim is an open-source software program purpose built for biomechanical modelling, simulation, and analysis. Data from the second half of the retraction phase and entire protraction phase (see Figure S13 below) for both the dominant and non-dominant legs were exported from Visual 3D to Opensim (v4.0) software. The musculoskeletal model was scaled to each individual using the subject's height, body mass and segment lengths (39). This model corresponds to the eight segments that were exported from Visual 3D. Simulations of muscle-tendon unit lengths were undertaken using the generic gait-2392 model within Opensim. The muscle-tendon unit (MTU) lengths of biceps femoris long head (BFlh), semimembranosus (SM), and semitendinosus (ST) were obtained for both DL and NDL in normal upright standing as well as non-fatigued and fatigued sprint running trials.

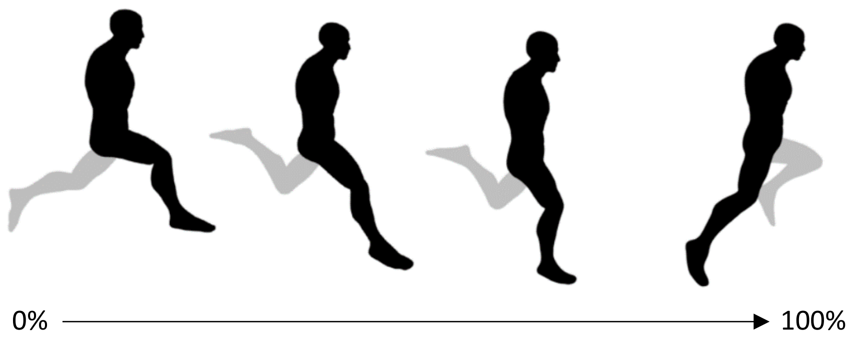


**Figure S13.** Second half of the retraction and throughout the entire protraction phase.

Peak MTU lengths were obtained from the second half of the retraction phase and entire protraction phase for BFlh, SM, and ST for both dominant and non-dominant legs in non-fatigued and fatigued sprint running trials. Peak MTU lengths were normalised to lengths during normal upright standing (34); therefore, values <1 can be interpreted as an increase in muscle-tendon unit length relative to normal upright standing.

Peak MTU lengths occurred between the second half of the retraction phase and the point just prior to foot-ground contact (i.e. late swing phase) for BFlh, SM, and ST in both limbs. No significant differences were observed in peak MTU lengths or the timing of peak MTU lengths between or within limbs during non-fatigued and fatigued sprinting, respectively. In fatigued sprinting, peak MTU lengths tended to decrease in DL but increase in NDL, although these changes did not reach statistical significance. In particular, BFlh (~1.3%) showed a greatest increase than SM and ST (see Figure S14). Any relation between inter-individual changes in MTU lengths and injury risk or running speed decline may be worthy of further investigation in future studies. However, our current findings are not consistent with the notion that an increase in anterior pelvic tilt directly increases hamstring MTU lengths.

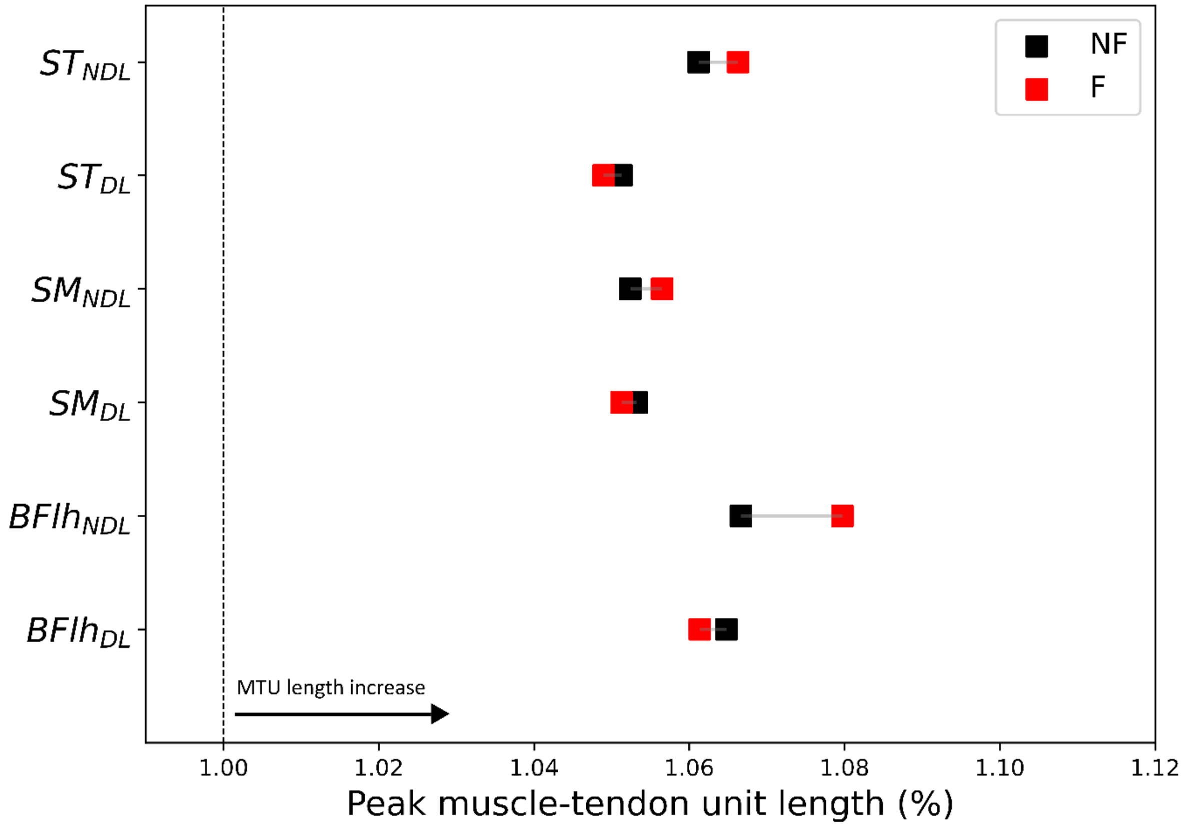


**Figure S14.** Comparison of peak muscle-tendon lengths in non-fatigued (NF: black squares) and fatigued (F: red squares) biceps Femoris long head ($BFlh_{DL}$ vs $BFlh_{NDL}$), semimembranosus ($SM_{DL}$ vs $SM_{NDL}$), and semitendinosus ($ST_{DL}$ vs $ST_{NDL}$) of the dominant (DL) and non-dominant legs (NDL) during sprinting. Vertical dashed line represents MTU length in normal upright standing. The Y axis displays the muscle followed by dominant (DL) or non-dominant leg (NDL). The X axis displays the peak muscle-tendon unit length; values < 1.00 indicate an increase in peak muscle-tendon unit length above normal upright standing.
